# Lipid chlorination mobilizes Neutrophil Elastase to initiate NETosis

**DOI:** 10.64898/2026.07.07.737103

**Authors:** Volker Brinkmann, Christian Goosman, Renate Krueger, Horst von Bernuth, Michael Mülleder, Kathrin Texoris-Taube, Brenda Moreira-Walsh, David A Ford, Alessandro Foti

## Abstract

Neutrophils are the first cells recruited to sites of infection, where they contain and kill pathogens through phagocytosis, degranulation, and the release of neutrophil extracellular traps (NETs) - web-like structures of decondensed chromatin that trap and destroy microbes. NET formation is essential for host defense, but when dysregulated it also contributes to disease, including sepsis, autoimmunity, and thrombosis. A central step in NET release is the translocation of neutrophil elastase (NE) from azurophilic granules to the nucleus, yet how NE crosses the granule membrane has remained unresolved. Using neutrophils and granules from healthy donors and from patients deficient in NADPH oxidase or myeloperoxidase (MPO), we show that MPO-derived hypochlorous acid chlorinates plasmalogens abundant in the granule membrane, generating 2-chlorofatty acids that permeabilize the granule and release NE into the cytosol. This reaction requires chloride, occurs within transient intracellular oxidant-rich compartments, and is both necessary and sufficient to trigger NE mobilization and NET formation. These findings identify lipid chlorination as the chemical link between the neutrophil oxidative burst and the execution of NETosis.

## Introduction

Neutrophils are the most abundant leukocytes in human blood and are rapidly recruited to sites of infection, where they provide immediate antimicrobial defense. They arrive primed for this task, carrying both preformed effector proteins and the enzymatic machinery required to generate a strong burst of reactive oxygen species (ROS) (*1*). These antimicrobial components are produced during neutrophil differentiation and stored in stage-specific granules, categorized as azurophilic, secondary, tertiary, and secretory vesicles, which permit regulated cargo delivery through membrane fusion (*2*). This cargo is deployed through three main antimicrobial strategies. In phagocytosis, pathogens are enclosed within membrane-bound phagosomes, where ROS and antimicrobial proteins cooperate to eliminate the ingested microbes, with granule cargo reaching the phagosome through fusion of granule and phagosomal membranes. Granules can also fuse directly with the plasma membrane, releasing their contents extracellularly through degranulation (*3*). Finally, neutrophils can extrude DNA-based fibers termed neutrophil extracellular traps (NETs) (*4*), web-like structures of decondensed chromatin decorated with granule and cytosolic proteins that trap and damage extracellular pathogens (*5*). NET formation is crucial for host defense: individuals and animal models carrying loss-of-function mutations in genes required for NET production show impaired clearance of, and increased susceptibility to, bacterial and fungal, alongside broader dysregulation of inflammatory responses (*6*). Yet NETs are a double-edged sword: when their production is excessive or their clearance is impaired, the same structures that protect the host instead become drivers of disease, contributing to the pathology of cystic fibrosis, preeclampsia, autoimmune disorders such as lupus, and vascular injury including thrombosis (*6*). Defining how NET release is triggered and controlled is therefore central to both infection biology and inflammatory disease.

NET release follows defined stimuli and proceeds through a specialized cell-death program, NETosis, in which chromatin decondenses and expands, the nuclear envelope breaks down, and the plasma membrane ultimately ruptures, expelling chromatin–protein complexes into the extracellular space (*7*). A central driver of this process is neutrophil elastase (NE), a serine protease normally confined to azurophilic granules, where it contributes to microbial killing within phagosomes. During NETosis, NE escapes its granule compartment and translocates to the nucleus, where it cleaves histones to relax chromatin (*8*). How NE exits across the granule membrane, however, remains one of the central unresolved questions in NETosis. NE release does not follow the rules of classical vesicle fusion, and ROS have long been implicated without a defined mechanism.

ROS generation is indispensable for neutrophil antimicrobial activity and for NETosis specifically: patients with chronic granulomatous disease (CGD), who lack functional NADPH oxidase, and individuals with complete myeloperoxidase (MPO) deficiency are highly prone to opportunistic infections, particularly fungal pathogens (*9*, *10*), and their neutrophils fail to generate NETs in response to physiological stimuli such as fungi or to the ROS-inducing mitogen phorbol myristate acetate (PMA) (*7*, *11*). Upon activation, NADPH oxidase produces superoxide, which is converted to hydrogen peroxide spontaneously or via superoxide dismutase 1 (SOD1) (*12*, *13*). MPO then uses hydrogen peroxide to generate hypohalous acids (HOX) and other oxidants that, beyond their cytotoxic role, can act as signaling molecules by chemically modifying target biomolecules (*14*, *15*). Importantly, how such short-lived oxidants exert spatially selective intracellular signaling during NETosis remains a central unanswered question. Although MPO is required for efficient NET formation, its mechanistic contribution is unclear (*8*, *16–19*). Additionally, 2-Chlorofatty acids metabolites of 2-chlorofatty aldehydes produced from MPO-derived hypochlorous acid (HOCl) targeting neutrophil plasmalogens has been implicated as a potential mediator of MPO caused NET formation (*20*).

Here, using primary human neutrophils and granules isolated from healthy donors and patients, we propose that ROS control NE mobilization through lipid chlorination. Formation of the 2-chlorofatty acids, 2-chloropalmitic acid and 2-chlorostearic acid after NET induction perturbs granule membrane integrity, allowing NE to enter the cytosol and translocate to the nucleus, thereby promoting NET release.

## Results

### NET formation depends on MPO-derived oxidants

Consistent with previous reports (*7*), stimulation of primary human neutrophils with the mitogen phorbol myristate acetate (PMA) or *Staphylococcus aureus* (MOI 20:1) induced robust NET release (Fig. 1A). In contrast, neutrophils from patients with chronic granulomatous disease (CGD) or complete myeloperoxidase (MPO) deficiency showed a marked reduction in NET formation (Fig. 1B–C), indicating that both NADPH oxidase–derived ROS and MPO activity are required. NET production was quantified by measuring extracellular DNA using the impermeant dye Sytox Green, which selectively labels DNA released into the extracellular space (Fig. 1D–F). As expected, CGD neutrophils failed to generate detectable ROS, whereas MPO-deficient neutrophils exhibited an enhanced but functionally altered oxidative burst (Fig. S1A). Because the primary product of NOX2 activity is hydrogen peroxide (HCOC), we asked whether other oxidants could substitute for endogenous ROS in promoting NETosis. Treatment with cumene hydroperoxide, potassium permanganate, or tert-butyl hydroperoxide did not induce NET release. In contrast, exogenous HCOC alone triggered substantial NET formation (Fig. S1B–C), suggesting a specific requirement for physiologically relevant ROS species. To determine whether the NET defect in CGD and MPO deficiency reflects the absence of HCOC or downstream hypohalous acids (HOX), we performed reconstitution experiments. Stimulation of healthy donor neutrophils with the mitogen PMA induced NET formation, and was blocked by inhibitors of ROS generation or serine proteases, confirming the dependence on both pathways (Fig. 1G). As expected, CGD neutrophils did not form NETs after mitogen stimulation. Addition of exogenous HCOC restored NET release, as did supplementation with the MPO product hypochlorous acid (NaOCl) (Fig. 1G). In MPO-deficient neutrophils, HCOC alone was insufficient to induce NETs, consistent with the lack of MPO activity. However, provision of NaOCl rescued partially NET formation in these cells (Fig. 1G). Together, these results indicate that MPO-derived oxidants act downstream of NOX2-generated HCOC to drive NETosis. We next asked whether NET formation requires a sustained oxidative burst or only an initial ROS signal. The oxidative burst occurs at the initial phase of NETosis and lasts for 60-70 minutes after activation, as shown by a luminol-based total ROS assay (Fig. 1H). Healthy neutrophils were stimulated with mitogen, and NOX2 activity was inhibited with diphenyleneiodonium (DPI) at defined time points after activation. Blocking ROS production at 30 minutes after stimulation abolished NET release, and inhibition at 45 minutes suppressed NET formation by approximately 50% (Fig. 1I). These data indicate that NETosis requires continued ROS production rather than a transient oxidative signal. Collectively, our findings demonstrate that MPO-derived oxidants are essential for NET formation with different stimuli and that efficient NETosis depends on sustained ROS generation.

**Figure 1.**
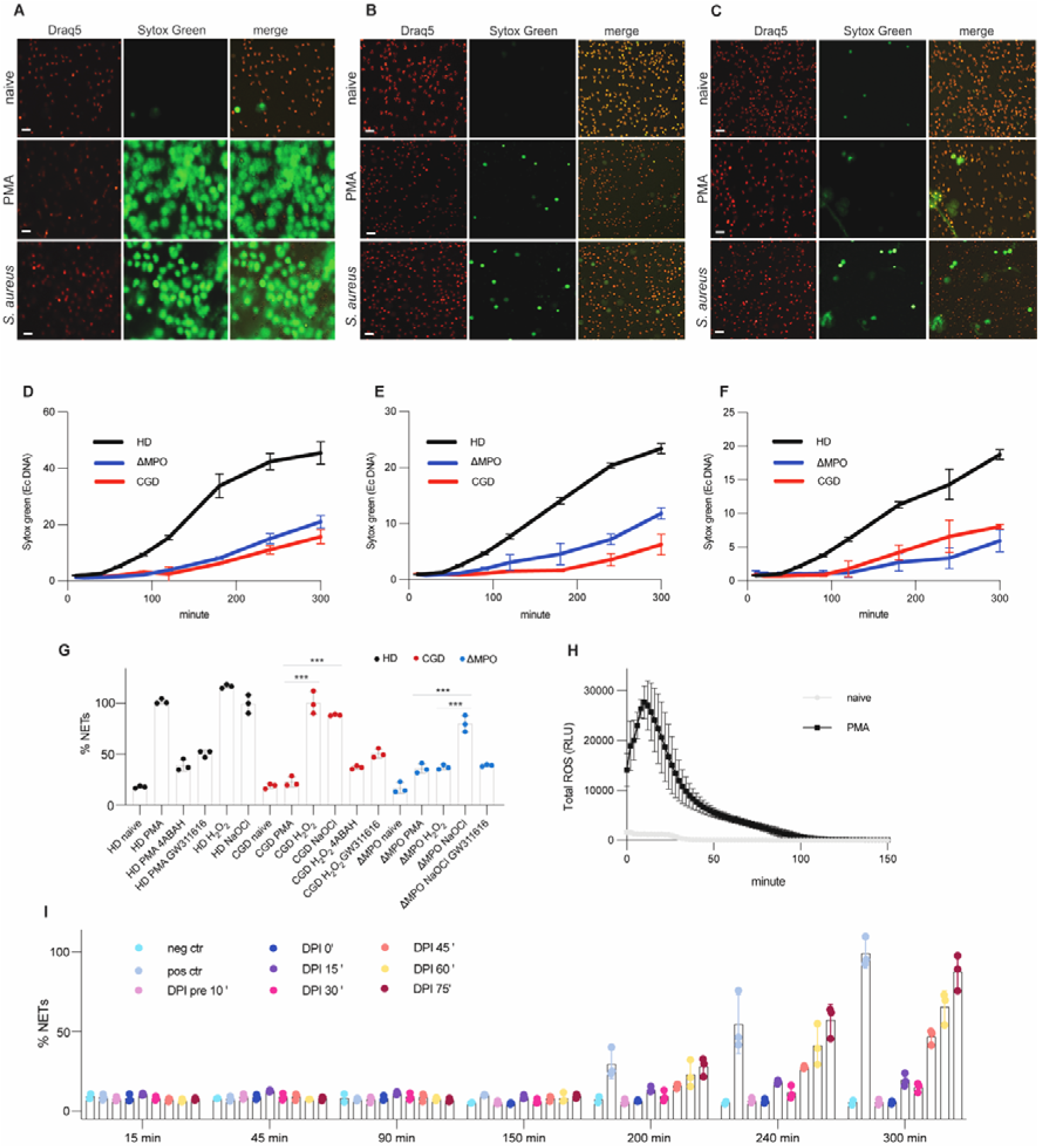
NET formation depends on MPO-derived oxidants. (A–C) Representative epifluorescence live microscopy images of neutrophils from a healthy donor (HD, N = 3) (A), a patient with chronic granulomatous disease (CGD, N = 3) (B), and a myeloperoxidase (MPO)-deficient patient (ΔMPO, N = 1) (C), stimulated for 5 h with PMA (100 nM) or infected with S. aureus (MOI 20:1). Draq5 stains intracellular DNA, and Sytox Green labels extracellular DNA. Scale bar, 20 μm. (D–F) Quantification and kinetics of NET formation in neutrophils from a healthy donor (D), a CGD patient (E), and an MPO-deficient patient (F) after PMA stimulation, based on Sytox Green fluorescence. Healthy donor (black), CGD patient (red), MPO-deficient patient (blue). Data represent means ± SEM. (G) Reconstitution and inhibition experiments in neutrophils from healthy donors (HD), CGD patients, and MPO-deficient patients stimulated with PMA, H□O□ (500 μM), or sodium hypochlorite (100 μM), with or without pretreatment with 4-ABAH (300 μM) or GW311616 (10 μM). Data points represent technical replicates. Healthy donors (N = 3), CGD patients (N = 3), MPO-deficient patient (N = 1). Data were analyzed by one-way ANOVA with Bonferroni multiple comparisons test; N=3. ***<0.001. (H) Total ROS production in healthy donor neutrophils after PMA stimulation (N = 3). (I) Quantification of NET formation in healthy donor neutrophils stimulated with PMA and treated with diphenyliodonium (1 μM) at different time points to inhibit ROS production (N = 3).

### Intracellular dynamics of MPO-derived oxidants during NETosis

To define the spatial and temporal dynamics of MPO-dependent oxidant production during NETosis, we used aminophenyl fluorescein (APF), a probe that selectively detects highly oxidizing MPO-derived species (*21*). Live epifluorescence imaging of primary human neutrophils stimulated with mitogen revealed the rapid appearance of discrete intracellular APF-positive compartments within ∼20 min of stimulation (Fig. 2A). These structures resembled membrane-bound vacuoles, as previously suggested (*7*), and we therefore termed them NETotic vacuoles. APF signal intensity increased during the first 30–60 min after stimulation and then declined, coincident with disappearance of the NETotic vacuoles. Thereafter, chromatin decondensation became apparent and extracellular NET release followed (Fig. S2A). To quantify these dynamics, we measured three independent parameters of APF signal in stimulated versus naive neutrophils over time: the number of individual APF-positive puncta per field, the number of discrete green foci clusters, and the total green fluorescent area per field. All three measures rose sharply after stimulation and peaked between 15 and 30 min (individual puncta per field, ∼26–32; green foci clusters, ∼22–25; total green area, ∼90–115 μm²), before declining rapidly by 60 min and returning to near-baseline levels by 120 min, while naive neutrophils remained APF-negative throughout the time course (Fig. 2B-D). This concordance across three orthogonal quantification metrics confirms that APF signal intensity increased during the first 20–30 min after stimulation and then declined, coincident with disappearance of the NETotic vacuoles, and establishes intracellular HOX accumulation as a discrete and self-limited early event of NETosis. Similar HOX dynamics were observed using live microbes. Infection of neutrophils with live *Candida albicans* induced intracellular HOX accumulation in a similar dynamic as mitogen stimulation (Fig. S2B). The HOX signal peaked at ∼45 min, diminished thereafter, and was followed by nuclear swelling and NET extrusion (Fig. S2B). These observations suggest that a transient intracellular burst of MPO-derived oxidants precedes and may license NET formation. To characterize the ultrastructural correlates of these events, we performed transmission electron microscopy (TEM) on mitogen-stimulated neutrophils from healthy donors, CGD patients, and DPI-treated controls. After 3 hours of stimulation, healthy neutrophils displayed extensive chromatin decondensation filling most of the cytoplasm, consistent with ongoing NETosis. In contrast, CGD neutrophils and DPI-treated cells retained large, intact NETotic vacuoles and failed to undergo chromatin expansion (Fig. S3A). In the absence of ROS production, these vacuoles appeared enlarged and persistent, suggesting that oxidant generation is required for their resolution.

**Figure 2.**
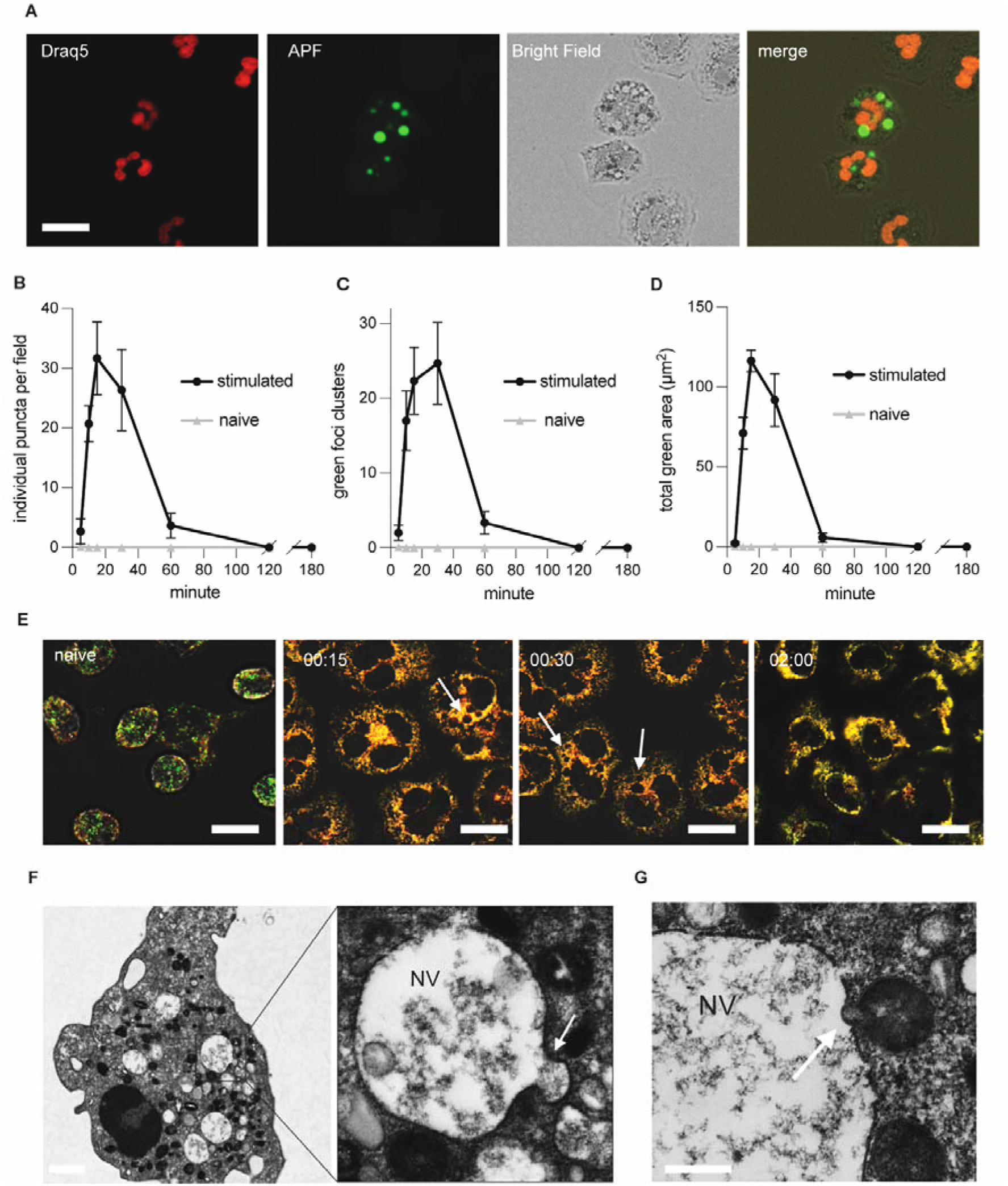
Intracellular dynamics of MPO-derived oxidants during NETosis. (A) Representative live-cell epifluorescence microscopy images of neutrophils from a healthy donor stimulated with PMA (100 nM) and imaged 20 min after activation. Draq5 stains intracellular DNA, and APF detects hypohalous acid production. Scale bar, 10 μm. (B) Quantification of individual APF□ puncta per field over time in neutrophils stimulated with PMA versus naïve controls. Data are mean ± SEM. (C) Quantification of the number of discrete green (APF□) foci/clusters per field over the same time course, comparing stimulated versus naïve neutrophils. Data are mean ± SEM. (D) Quantification of total APF□ (green) fluorescent area (μm²) per field over time, comparing stimulated versus naïve neutrophils. Data are mean ± SEM. (E) Confocal immunofluorescence images of healthy donor neutrophils left untreated (naïve) or stimulated with PMA for 15, 30, or 120 min and stained for MPO and gp91^phox (N = 3). Arrows indicate sites of MPO/gp91^phox co-localization at the NETotic vacuole (NV) membrane. Scale bar, 5 μm. (F) Transmission electron microscopy of a PMA-stimulated neutrophil, with inset showing higher magnification of the NV; arrow indicates a granule in apposition to the NV membrane. Scale bar, 1 μm. (G) Transmission electron micrograph of NV, with arrow indicating a granule fusing with the NV membrane. Scale bar, 100 nm.

Because MPO resides in azurophilic granules whereas the NADPH oxidase complex localizes to secondary granules, we hypothesized that formation of HOX-rich vacuoles requires fusion between distinct granule populations. Immunofluorescence analysis of mitogen-stimulated neutrophils showed colocalization of MPO with gp91^phox, a core NADPH oxidase component, around NETotic vacuoles (Fig. 2E). Similar colocalization was observed for neutrophil elastase and p22^phox (Fig. S3B), supporting the idea that granule mixing precedes HOX generation. We performed additional TEM analysis of stimulated neutrophils and observed electron-dense granules apposed to and fusing with the limiting membrane of NETotic vacuoles, capturing sequential stages of granule-vacuole fusion — docking, membrane apposition, and merging (arrowheads, Fig. 2F-G; Fig. S4A). Interestingly, this process recalls a similar mechanism observed during phagocytosis (*22*).

Together, these data indicate that NETosis involves the transient formation of intracellular HOX-enriched NETotic vacuoles generated through granule fusion. The subsequent disappearance of these compartments correlates with chromatin decondensation and NET release, suggesting that resolution of NETotic vacuoles is a prerequisite for execution of the NETotic program.

### Dynamic behaviour of NETotic vacuoles and MPO-dependent neutrophil elastase mobilization

To better define the formation and resolution of NETotic vacuoles, we performed three-dimensional live confocal imaging of APF-stained neutrophils after stimulation. NETotic vacuoles behaved as highly dynamic intracellular compartments. They moved rapidly within the cytoplasm and displayed marked changes in size and number over time (Fig. 3A). Consistent with our earlier observations, dissolution of these vacuoles preceded chromatin decondensation and NET release, indicating that their resolution is a prerequisite for NETosis. Given the apparent role of granule remodeling in this process, we next isolated neutrophil granules using Percoll density fractionation (*23*). This approach allowed separation of distinct granule populations, which were validated by immunoblotting for established markers of azurophilic, secondary, tertiary granules and secretory vescicles (Fig. 3B). Label-free quantitative proteomics further confirmed the expected protein composition of each fraction (Fig. S5A-D). Ultrastructural analysis by scanning electron microscopy (SEM) showed that azurophilic granules have diameters of approximately 150–250 nm (Fig. 3C; Fig. S6A–C). Immunogold labeling verified the presence of NE within these organelles (Fig. 3D). We then asked how NE exits azurophilic granules during NETosis. Purified azurophilic granules were incubated with a fluorogenic elastase substrate (elastin) that cannot cross intact granule membranes. Cleavage of the substrate generates fluorescence and therefore reports NE release into the surrounding medium. As a positive control, detergent solubilization of granules with NP-40 produced maximal substrate cleavage (Fig. 3E). Treatment with HCOC induced a time-dependent increase in NE activity outside the granules, reaching a peak at ∼60 min (Fig. 3E). This effect was blocked by the MPO inhibitor 4-aminobenzoic acid hydrazide (4-ABAH), indicating that NE mobilization requires MPO activity. A dose–response analysis confirmed that NE release scaled with HCOC concentration (Fig. 3F). Similar results were obtained using alternative NE substrates, including fluorescent streptavidin, and were influenced by pH (Fig. S7A–D). We also confirmed released of NE translocation upon H_2_O_2_ stimulation by western blot (Fig. 3G). Additionally, transmission electron microscopy analysis revealed that H_2_O_2_ partially altered the ultrastructure of the granules (Fig. S8A-B). To directly test the requirement for MPO, we purified azurophilic granules from MPO-deficient patients. In contrast to granules from healthy donors, MPO-deficient granules failed to release NE in response to HCOC (Fig. 3H). Independent assays confirmed the absence of MPO activity from these granules (Fig. 3I). Together, these experiments demonstrate that NE mobilization from azurophilic granules is driven by MPO-dependent enzymatic activity. Loss of MPO activity abolishes NE release, supporting a model in which HCOC–MPO chemistry destabilizes granule membranes to permit NE translocation, a key step in NETotic progression.

**Figure 3.**
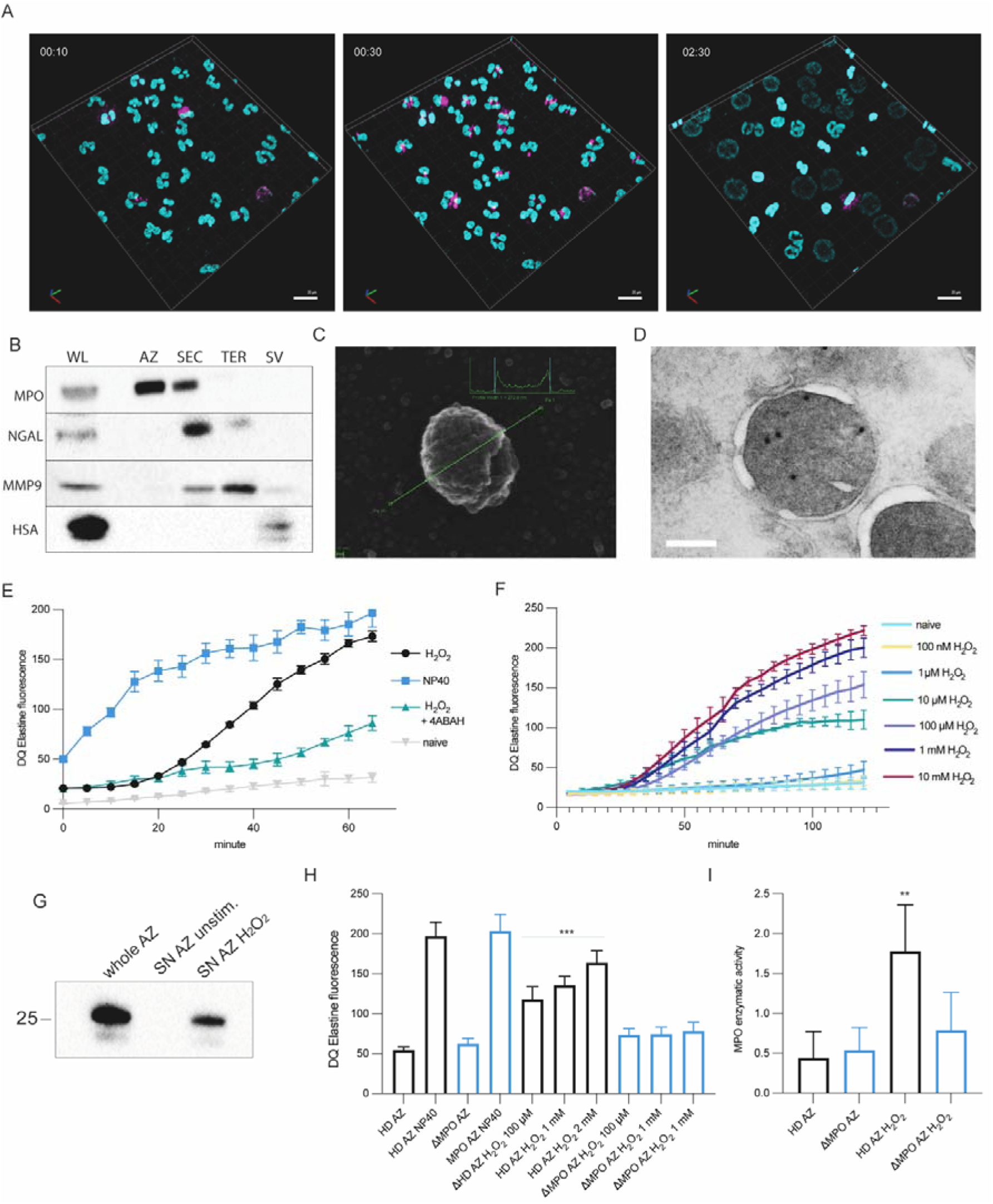
Dynamic behaviour of NETotic vacuoles and MPO-dependent neutrophil elastase mobilization. (A) Time-course three-dimensional live confocal microscopy images of neutrophils from a healthy donor stimulated with PMA (100 nM) and imaged at 10, 30, and 150 min after stimulation (N = 3). Light blue fluorescence indicates intracellular DNA, and magenta fluorescence indicates intracellular hypohalous acid formation. Scale bar, 20 μm. (B) Western blot analysis of purified neutrophil granule fractions from healthy donors. Azurophilic granules (AZ) were probed for MPO, secondary granules (SEC) for NGAL, tertiary granules (TER) for MMP9, and secretory vesicles (SV) for HSA (human serum albumin). WL, whole-cell lysate (N = 5). (C) Scanning electron microscopy image of a purified azurophilic granule from a healthy donor (N = 3). (D) Transmission electron microscopy image of purified azurophilic granules immunogold-labeled for NE (N = 4). Scale bar, 200 nm. (E) Neutrophil elastase (NE) translocation assay. NE activity was measured by DQ-elastin fluorescence over time in purified azurophilic granules treated with H□O□ (100 μM) in the presence or absence of the MPO inhibitor 4-ABAH (300 μM), lysed with NP-40 (positive control), or left untreated (naïve, negative control) (N = 5). Data are mean ± SEM. (F) NE activity (DQ-elastin fluorescence) in azurophilic granules treated with increasing concentrations of H□O□ (100 nM–10 mM) over time, compared with naïve (untreated) granules (N = 3). Data are mean ± SEM. (G) Western blot detecting NE release from purified azurophilic granules. Whole granule lysate (whole AZ) is compared with the granule supernatant (SN) under unstimulated conditions or after treatment with H□O□, showing translocation of NE (∼25 kDa) into the supernatant. (H) Quantification of NE activity (DQ-elastin fluorescence, 60 min) in azurophilic granules from healthy donors (HD AZ, N=*3*) and MPO-deficient patients (ΔMPO AZ, N=*1*), untreated, treated with NP-40 (positive control), or treated with increasing concentrations of H□O□ (100 μM–2 mM). Data were analyzed by one-way ANOVA with Bonferroni multiple comparisons test; N=3. ***P < 0.001. Data are mean ± SEM. (I) MPO enzymatic activity in azurophilic granules from healthy donors (HD AZ, N=*3*) and MPO-deficient patient (ΔMPO AZ, N=*1*), untreated or treated with H□O□. Data were analyzed by one-way ANOVA with Bonferroni multiple comparisons test; **P < 0.01. Data are mean ± SEM.

### Chloride-dependent MPO activity drives neutrophil elastase mobilization and NET formation

Our results indicate that MPO catalytic activity is required for NE release from azurophilic granules. MPO, however, can utilize several substrates, and under physiological conditions chloride is the predominant anion available, leading to the production of hypochlorous acid (HOCl) as the major oxidant. We therefore asked whether chloride availability limits NET formation. Primary neutrophils from healthy donors were incubated in a customized chloride-free medium in which chloride was replaced with gluconate , and NET release was quantified after stimulation with mitogen. Removal of extracellular chloride markedly reduced NET formation, whereas re-addition of chloride restored NET production in a dose-dependent manner (Fig. 4A–B). We observed comparable NETs reduction also after stimulating neutrophils with *S. aureus* or *C. albicans* (Fig. S9A-B). These results indicate that chloride availability is required for efficient NETosis. To test whether chloride is also necessary for NE mobilization, we examined purified azurophilic granules. Granules incubated in chloride-free medium failed to release NE upon HCOC treatment. Reconstitution with physiological chloride concentrations restored NE translocation (Fig. 4C). Importantly, the extent of NE release closely correlated with MPO chlorination activity measured in a parallel enzymatic assay (Fig. 4D), suggesting that MPO-mediated chlorination reaction drives this process. To determine whether chlorination is sufficient to trigger NE mobilization, we exposed purified granules to defined chlorinating agents, including hypochlorite, monochloramines, and dichloramines. Each reagent induced robust NE release from azurophilic granules (Fig. 4E), supporting a direct role for chlorinating oxidants in destabilizing granule membranes. Together, these findings demonstrate that chloride-dependent MPO activity is essential for both NE translocation and NET formation.

**Figure 4.**
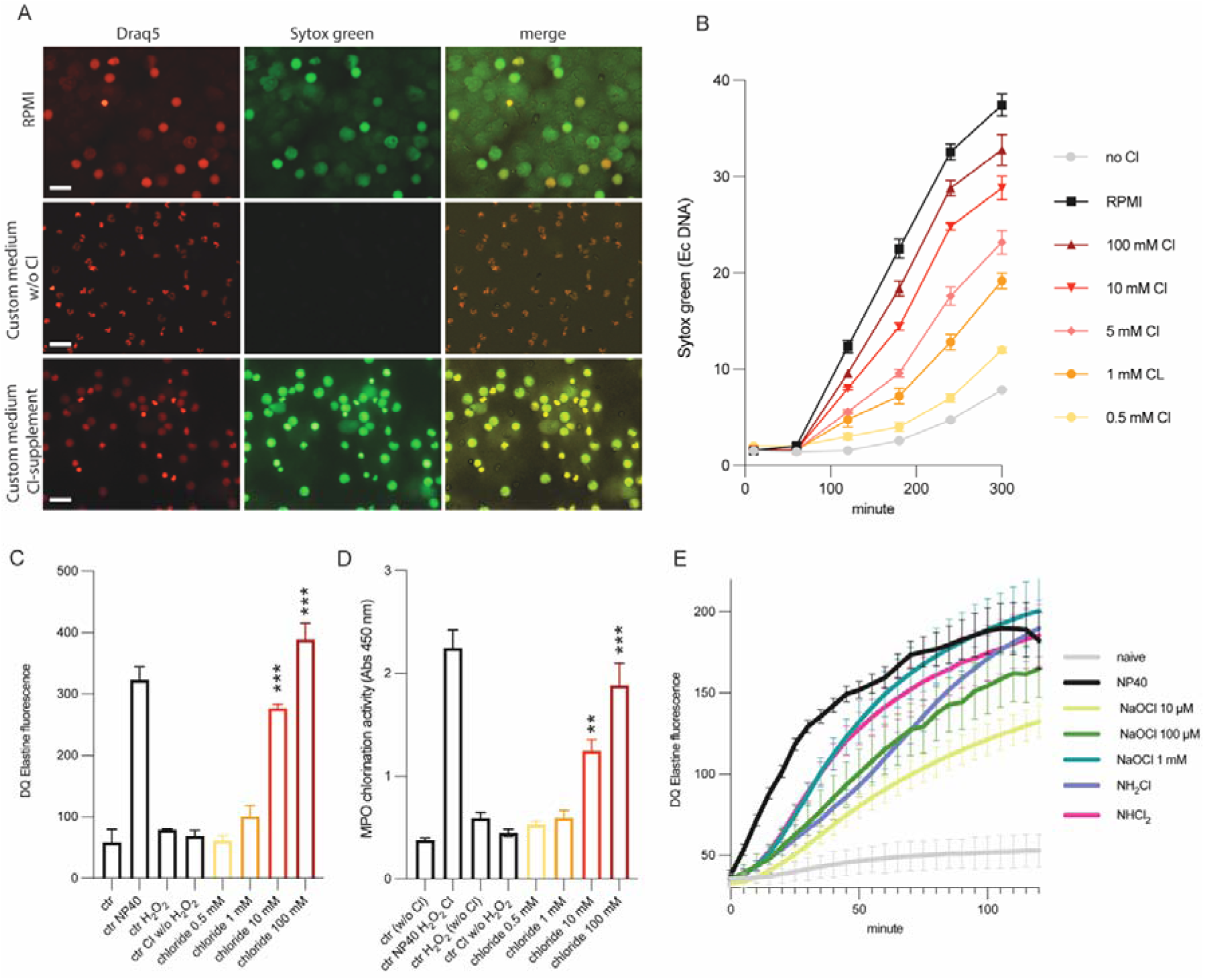
Chloride-dependent MPO activity drives neutrophil elastase mobilization and NET formation. (A) Representative live-cell epifluorescence microscopy images of neutrophils from a healthy donor stimulated for 5 h with PMA (100 nM) in standard RPMI medium, chloride-free medium, or chloride-restored medium (N = 5). Draq5 stains intracellular DNA, and Sytox Green labels extracellular DNA. Scale bar, 20 μm. (B) Quantification and kinetics of NET formation in neutrophils from healthy donors cultured in standard RPMI medium, chloride-free medium, or chloride-restored medium at increasing chloride concentrations after PMA stimulation, based on Sytox Green fluorescence (N = 5). Data are mean ± SEM. (C) Neutrophil elastase (NE) translocation assay in azurophilic granules from healthy donors. NE activity was measured by DQ-elastin fluorescence 60 min after treatment with H□O□ (100 μM) in chloride-free medium or chloride-restored medium at increasing chloride concentrations. NP-40 was used as a positive control, and untreated samples served as negative controls (N = 4). Data were analyzed by one-way ANOVA with Bonferroni multiple comparisons test; N=3. ***P < 0.001. Data are mean ± SEM. (D) MPO enzymatic activity measured in azurophilic granules from healthy donors incubated in chloride-free medium or chloride-restored medium at increasing chloride concentrations after H□O□ stimulation (N = 4). Data were analyzed by one-way ANOVA with Bonferroni multiple comparisons test; N=3. **P < 0.01. Data are mean ± SEM. (E) NE activity measured by DQ-elastin fluorescence in granules treated with chlorinated oxidants, including sodium hypochlorite (increasing concentrations), monochloramine (100 μM), and dichloramine (100 μM), monitored over time (N = 3). Data are mean ± SEM.

### Granule plasmalogens constitute a substrate reservoir for MPO**D**dependent lipid chlorination

Previous studies reported that stimulated neutrophils accumulate chlorinated lipids derived from plasmalogens and that these species can induce NET formation (*24*). Because lipid chlorination requires a plasmalogen substrate positioned within reach of MPO (*25*), we first defined the phospholipid landscape of the neutrophil granules. We resolved the four granule populations — azurophilic, secondary, tertiary granules and secretory vesicles — by Percoll fractionation and quantified their phosphatidylcholine (PC) and phosphatidylethanolamine (PE) species by liquid chromatography–mass spectrometry (Fig. 5A-B). Both headCgroup classes were detected across all four compartments, and their molecularCspecies profiles were remarkably conserved between subtypes; the granule populations differed principally in absolute lipid load, which increased progressively from azurophilic granules to secretory vesicles, rather than in composition. When species were grouped by their snC1 linkage, PC comprised a mixture of diacyl (∼65%), alkylCacyl (∼20%) and plasmalogen (∼15%) species (Fig. 5C). In striking contrast, PE was composed almost entirely of plasmalogens, which exceeded 92% of all PE species in every compartment and reached ∼96% in azurophilic granules (Fig. 5D). Thus the vinylCether plasmalogens that serve as the substrate for hypochlorous acid are the dominant ethanolamine phospholipid of neutrophil granules, and they are abundant in the azurophilic compartment that also harbours MPO — coClocalizing enzyme and substrate within the same organelle.

**Figure 5.**
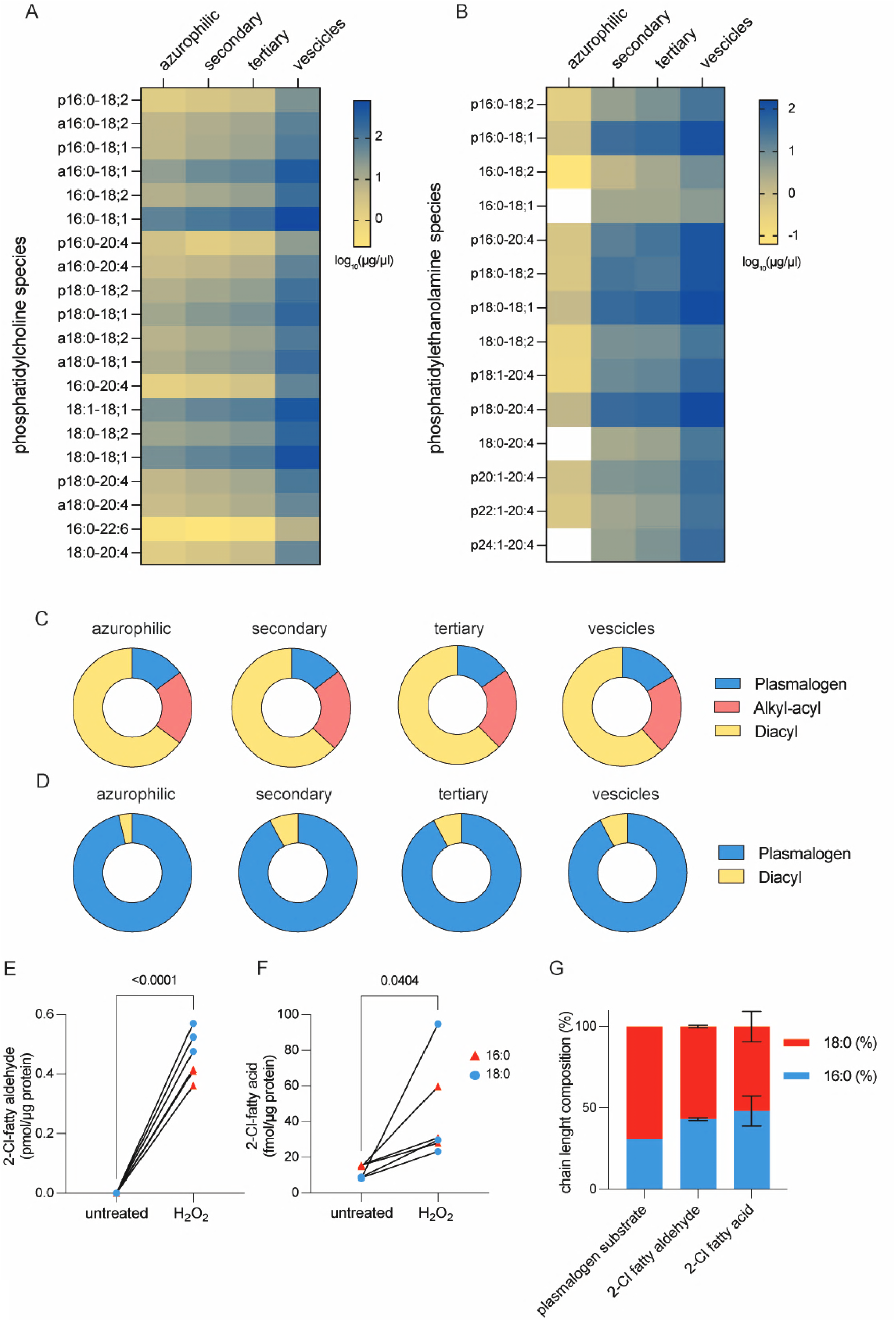
The neutrophil granule phospholipidome is plasmalogen□rich and supports MPO□dependent generation of chlorinated lipids. (A) Heatmap of phosphatidylcholine (PC) molecular species (rows) across four purified granule subtypes: azurophilic, secondary, tertiary granules and secretory vesicles (columns). Colour encodes abundance as log□□(µg/µl). Species are annotated by sn□1 linkage: p, plasmalogen (1□O□alk□1′□enyl/vinyl□ether); a, alkyl□acyl (1□O□alkyl); no prefix, diacyl. Values are means of n = 5 azurophilic, 3 secondary/tertiary/vesicles independent preparations. (B) As in (A) for phosphatidylethanolamine (PE) species. White cells denote species not detected. Values are means of n = 5 azurophilic, 3 secondary/tertiary/vesicles independent preparations. (C) Relative class composition of PC — plasmalogen, alkyl□acyl and diacyl — in each granule subtype (same preparations as A). (D) Relative class composition of PE — plasmalogen and diacyl — in each granule subtype (same preparations as B). PE is composed almost entirely of plasmalogens (∼92–96%) across all subtypes. (E) 2□Chlorofatty aldehyde content of isolated granules, untreated versus after treatment with H□O□ (100 µM). Red triangles, 16:0 (2□chlorohexadecanal/2□chloropalmitaldehyde); blue circles, 18:0 (2□chlorostearaldehyde). Lines connect paired measurements from the same preparation; values are pmol per µg protein (n = 3). The P value (untreated vs H□O□) is indicated. (F) 2□Chlorofatty acid content of the same granules, untreated versus H□O□ (100 µM). Red triangles, 16:0 (2□chloropalmitic acid); blue circles, 18:0 (2□chlorostearic acid). Values are fmol per µg protein (n = 3). The P value is indicated. (G) Acyl□chain composition (% 16:0 versus 18:0) of the granule plasmalogen substrate pool, the 2□chlorofatty aldehyde product and the 2□chlorofatty acid product. The plasmalogen substrate is ∼31% 16:0 / 69% 18:0, whereas both chlorinated products are enriched in 16:0 (aldehyde 43 ± 1%; acid 48 ± 9% 16:0), indicating preferential generation of the 16:0 (palmitoyl) chlorolipid relative to its abundance in the substrate. Bars are mean ± SD (substrate, composite value; products n = 3).

We next asked whether activation of granuleCresident MPO is sufficient to chlorinate this endogenous plasmalogen pool. HOCl attacks the *sn*C1 vinylCether bond of plasmalogens to release a 2Cchlorofatty aldehyde, which can be further oxidized to the corresponding 2Cchlorofatty acid. We therefore incubated isolated granules with hydrogen peroxide (HCOC, 100 µM), the coCsubstrate of MPO, and measured both chlorinated species. Under resting conditions, 2Cchlorofatty aldehydes were essentially undetectable; addition of HCOC triggered their robust accumulation, generating both the 16:0 (2Cchloropalmitaldehyde) and 18:0 (2Cchlorostearaldehyde) species (Fig. 5E). The downstream 2Cchlorofatty acids, 2Cchloropalmitic and 2Cchlorostearic acid, were present at low levels in untreated granules and rose significantly upon HCOC treatment (Fig. 5F). Because these reactions occurred in purified granules supplied only with HCOC, they demonstrate that the chlorinating machinery, its peroxide substrate and the plasmalogen target are all contained within the granule itself, enabling chlorinated lipids to be produced *in situ*.

Finally, we tested whether the chlorinated products could be traced to the granule plasmalogen pool from which they are predicted to arise. Because the 2Cchlorofatty aldehyde derives directly from the *sn*C1 vinylCether chain, we compared the 16:0/18:0 composition of the granule plasmalogen substrate with that of the chlorinated products (Fig. 5G). The plasmalogen pool was dominated by 18:0 *sn*C1 chains (∼69%, versus ∼31% 16:0), a distribution that was essentially invariant across granule subtypes. Both chlorinated products mirrored this pool in being derived from saturated snC1 chains, but were consistently and disproportionately enriched in the 16:0 species relative to the substrate (16:0 fraction: aldehyde 43 ± 1%; acid 48 ± 9%). This preferential generation of the 2-chloropalmitaldehyde provides a substrateClevel explanation for the prominence of 2Cchloropalmitic acid among neutrophil chlorinated lipids and links the chlorinated products quantitatively to the granule plasmalogen reservoir. Together, these data establish that neutrophil granules are a plasmalogenCrich compartment in which MPO activity generates 2Cchlorofatty aldehydes and acids *in situ*, defining the chemical step that connects the oxidative burst to the chlorinated lipids formation.

### Chlorinated plasmalogen-derived lipids promote NET formation and neutrophil elastase release

To validate and extend chlorinated the previous finding suggesting that lipids derived from plasmalogens can induce NET formation (*24*), we stimulated primary human neutrophils with palmitic acid or 2-chloropalmitic acid, using the mitogen as a positive control. Whereas palmitic acid had no detectable effect, 2-chloropalmitic acid induced robust NET release comparable to mitogen stimulation (Fig. 6A). We next examined structurally related lipids, including stearic acid, 2-chlorostearic acid, myristic acid, and arachidic acid. Among these, only 2-chlorostearic acid triggered NET formation, and with lower potency than 2-chloropalmitic acid (Fig. 6B), indicating a selective activity of chlorinated saturated fatty acids. NET induction by 2-chloropalmitic acid was blocked by serine protease inhibitors (Fig. 6C), placing its action upstream of neutrophil elastase (NE) activation during NETosis. We therefore hypothesized that chlorinated palmitic acid may link MPO-derived oxidant production to NE mobilization. To test this hypothesis, purified azurophilic granules were incubated with increasing concentrations of 2-chloropalmitic acid. This treatment induced a strong and dose-dependent release of NE from granules, whereas palmitic acid had no effect (Fig. 6D). Additional lipid species were tested in parallel, confirming that 2-chloropalmitic acid was the most potent inducer of NE translocation (Fig. 6E). Additionally, we confirmed 2-chloropalmitic acid-dependent translocation of NE by western blot (Fig. 6F). To determine whether chlorinated lipids directly affect granule membrane integrity, purified granules were incubated with 2-chloropalmitic acid and then separated into pellet and supernatant fractions by ultracentrifugation. SDS–PAGE analysis revealed a progressive loss of protein content from the granule pellet with increasing concentrations of chlorinated lipid, accompanied by a corresponding increase in proteins detected in the supernatant (Fig. S10). These results indicate that chlorinated palmitic acid perturbs granule membranes and promotes leakage of luminal proteins. Together, these findings identify 2-chloropalmitic acid and 2-chlorostearic acid as key effectors downstream of MPO activity. By perturbing azurophilic granule membrane integrity, this chlorinated lipid facilitates NE release into the cytosol and thereby promotes NET formation. This mechanism provides a direct biochemical link between MPO-driven lipid chlorination and the execution of NETosis.

**Figure 6.**
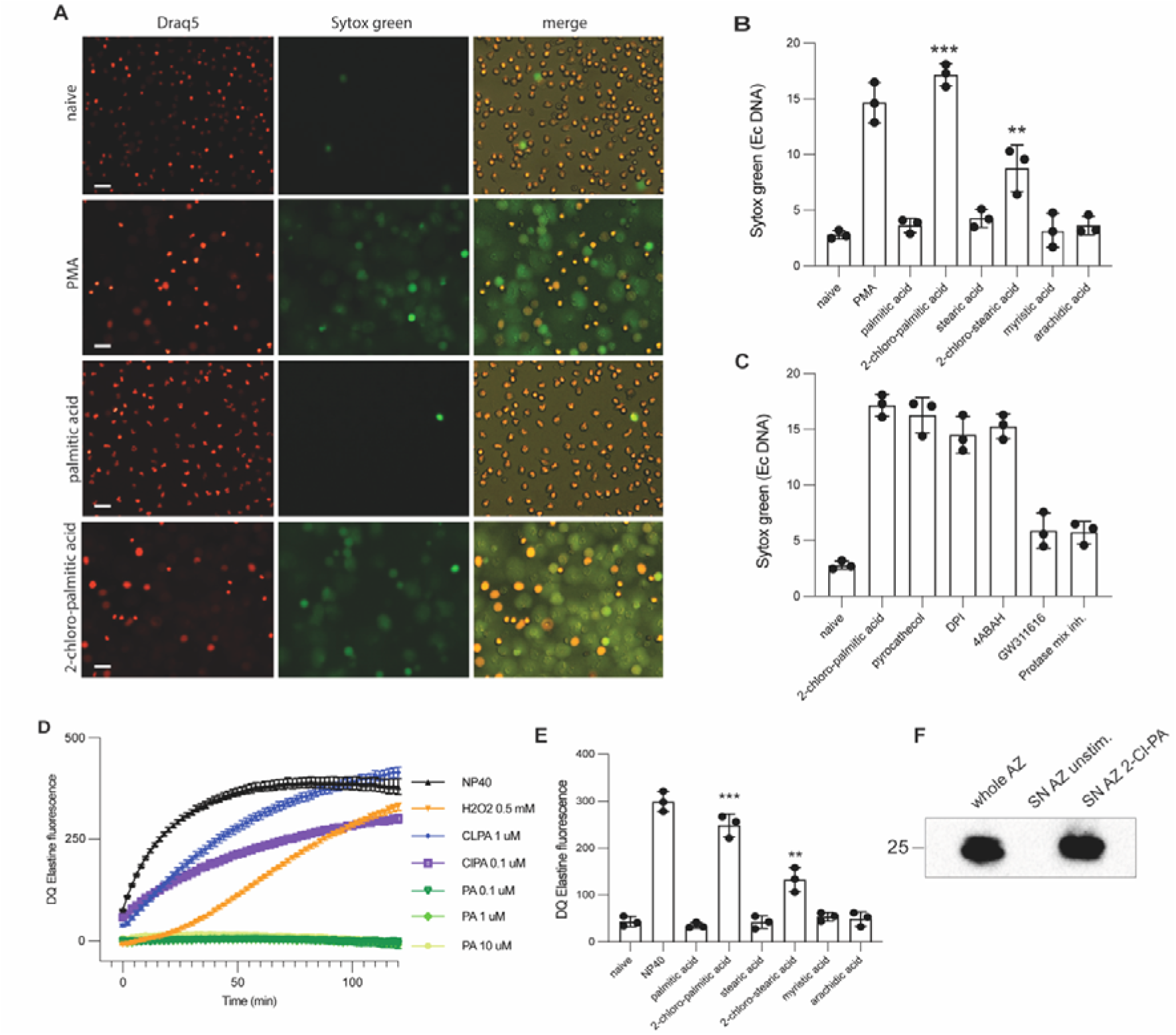
Chlorinated plasmalogen-derived lipids promote NET formation and neutrophil elastase release. (A) Representative live-cell epifluorescence microscopy images of neutrophils from healthy donors stimulated for 5 h with PMA (100 nM), palmitic acid (10 μM), or 2-chloropalmitic acid (10 μM), or left untreated (naïve) (N = 3). Draq5 stains intracellular DNA, and Sytox Green labels extracellular DNA. Scale bar, 20 μm. (B) Quantification of NET formation in healthy donor (HD) neutrophils stimulated with PMA or the indicated fatty acids (10 μM), based on Sytox Green fluorescence (N = 3). Data were analyzed by one-way ANOVA with Bonferroni multiple comparisons test; N=3. Data are mean ± SEM.**P < 0.01, ***P < 0.001 vs. naïve. (C) Quantification of NET formation in healthy donor neutrophils stimulated with 2-chloropalmitic acid (10 μM) following pretreatment with the indicated inhibitors — pyrocatechol, DPI, 4-ABAH, GW311616, or a protease mixture inhibitor — based on Sytox Green fluorescence (N = 3). Data are mean ± SEM. (D) Neutrophil elastase (NE) activity, measured by DQ-elastin fluorescence over time, in azurophilic granules treated with NP-40 (positive control), H□O□ (0.5 mM), 2-chloropalmitic acid (0.1 or 1 μM), or palmitic acid (0.1, 1, or 10 μM) (N = 3). Data are mean ± SEM. (E) Quantification of NE activity (DQ-elastin fluorescence) in azurophilic granules treated with NP-40 (positive control) or the indicated fatty acids, compared with untreated (naïve) granules (N = 3). Data were analyzed by one-way ANOVA with Bonferroni multiple comparisons test; N=3. Data are mean ± SEM.**P < 0.01, ***P < 0.001 vs. naïve (F) Western blot for neutrophil elastase (NE, ∼25 kDa) in whole azurophilic granule lysate (whole AZ) and in the granule supernatant (SN) under unstimulated conditions or after treatment with 2-chloropalmitic acid, showing NE release into the supernatant (N = 3).

## Discussion

Neutrophils are indispensable effector cells of innate immunity, deploying a coordinated repertoire of oxidative and nonCoxidative antimicrobial mechanisms. Among these is the formation of neutrophil extracellular traps (NETs), networks of decondensed chromatin bound to granule and cytosolic proteins that immobilize and kill microbes. NETs are released through NETosis, a regulated cell death in which chromatin decondenses and expands, the nuclear envelope disassembles, and the plasma membrane ruptures to expel the chromatin. The same process that contains pathogens drives tissue injury when dysregulated, contributing to multiple pathological conditions such as cystic fibrosis, sepsis, thrombosis and autoimmunity (*3*). A central event in NETosis is the mobilization of neutrophil elastase (NE) from azurophilic granules to the nucleus, where it processes histones to permit chromatin decondensation. How NE exits the granule has remained unclear, as it does not rely on the classical membrane fusion that delivers granule cargo to the phagosome. In this study, we identify a novel mechanism that couples the neutrophil oxidative burst to NE mobilization through the chemical modification of granule lipids, revealing an intersection between MPOCderived oxidant chemistry and the execution of NETosis.

Hypohalous acids are typically regarded as broadly cytotoxic effectors (*13*). Increasingly, however, MPOCderived oxidants are recognized to react selectively with neutrophil components, modifying host molecules rather than indiscriminately degrading them (*15*, *26*, *27*). Our findings extend this principle from proteins to membrane lipids. Upon activation, MPO uses hydrogen peroxide and chloride to generate hypochlorous acid, which attacks the *sn*C1 vinylCether bond of plasmalogens to yield 2Cchlorofatty aldehydes and their oxidation products, 2Cchlorofatty acids (*24*). Here we present the first, to our knowledge, lipidomic landscape of the different neutrophil granules isolated from healthy donor human neutrophils (*20*, *25*, *28*). MPO is stored in neutrophil granule together with an abundant plasmalogen substrate, and granule phosphatidylethanolamine consists almost entirely of plasmalogens. Isolated granules supplied with hydrogen peroxide generated both chlorinated species *in situ*, with the aldehyde predominating, and the chainClength distribution of the products reflected that of the plasmalogen pool, linking the chlorinated lipids to their granule source.

Through complementary cellular, biochemical and lipidomic analyses, we show that these chlorinated lipids accompany the release of NE from azurophilic granules. Hydrogen peroxide triggered NE mobilization from isolated granules, an effect abolished by the MPO inhibitor 4CABAH and absent in granules from MPOCdeficient patients, establishing a requirement for MPO catalytic activity. Defined chlorinating agents reproduced this effect, and exogenous chlorinated fatty acids were sufficient to perturb granule membrane integrity. In intact neutrophils, MPOCderived oxidants accumulated transiently within discrete intracellular compartments, which we term NETotic vacuoles, formed at sites of convergence between MPOC and NADPH oxidase–bearing granules; their resolution preceded chromatin decondensation and NET release. Exogenous chlorinated fatty acids were sufficient to induce NETs and the process was inhibited from serine protease inhibitors, further confirming the axis MPO-chlorinated lipids-NE activity. Together, these observations support a model in which MPOCdriven plasmalogen chlorination destabilizes the azurophilic granule membrane, releasing NE to the cytosol and licensing its translocation to the nucleus. The chloride dependence of this process underscores its enzymatic basis. Chloride is the predominant physiological substrate of MPO, and its removal abolished both NE release from isolated granules and NET formation in cells, each restored by chloride in proportion to MPO chlorination activity.

Chlorinated lipids derived from plasmalogens are chemically stable and accumulate in neutrophil and monocytes-rich inflammation (*29*, *30*, *31*, *32*), positioning them as important candidate markers of MPO activity and NETCassociated pathology in diseases such as sepsis (*33*, *34*, *35*). More fundamentally, our findings indicate that the oxidative output of MPO is a key signal during NETosis through a discrete lipid modification.

In conclusion, our study identifies MPOCdriven lipid chlorination as a step that connects the neutrophil oxidative burst to the mobilization of neutrophil elastase during NETosis. By chemically destabilizing the granule membrane that confines NE, MPOCderived oxidants couple oxidant production to the proteolytic NE activity that drive chromatin decondensation and NET release. This mechanism reveals a previously unrecognized chemical layer linking hypochlorous acid signalling with the execution of NETosis.

## Materials and Methods

### Chemicals

All reagents were purchased from common vendors of laboratory reagents, such as Sigma-Aldrich, Thermo Fisher Scientific or VWR Deutschland, unless otherwise stated.

### Isolation of human neutrophils

The ethics council of the Charité – Universitätsmedizin Berlin (Germany) approved blood sampling, and all donors gave written informed consent according to the Declaration of Helsinki. Human neutrophils were isolated by a two-step density separation as described previously (*36*). Briefly, freshly drawn peripheral blood was layered on an equal volume of Histopaque 1119 (Sigma-Aldrich) and centrifuged at 800 g for 20 min. The peripheral blood mononuclear cell and neutrophil layers were collected separately, washed with PBS containing 0.2% human serum albumin (HSA) and pelleted at 300 g for 10 min. The neutrophil pellet was resuspended in PBS/0.2% HSA, layered on a discontinuous Percoll gradient (85% to 65% in 2-ml layers) and centrifuged at 800 g for 20 min. The neutrophil-containing band was collected, washed in PBS/0.2% HSA and pelleted for 10 min at 300 g. Cell number was determined with a CASY cell counter (OMNI Life Science). Neutrophils were resuspended in RPMI-1640 (Gibco) supplemented with 5 mM HEPES and 0.2% HSA and used immediately.

### Patients

Blood samples were collected according to the Declaration of Helsinki with written informed consent. Samples from patients with chronic granulomatous disease (CGD) and from the myeloperoxidase (MPO)-deficient donor were collected with approval from the ethics committee of the Charité – Universitätsmedizin Berlin. Patients with X-linked CGD harboured mutations in the NADPH-oxidase subunit gene CYBB (e.g. c.742dupA; c.868C>T), and the MPO-deficient donor carried a homozygous splice-site mutation (c.2031-2A>C) generating null alleles and a catalytically inactive, immature protein. The number of independent donors or patients analysed in each experiment is stated in the corresponding figure legend

### Microbial culture

*Staphylococcus aureus* strain USA300 was inoculated into tryptic soy broth (TSB) and grown at 37 °C for 18 h with shaking; the culture was then passaged into fresh TSB and grown for ∼2 h to mid-logarithmic phase, harvested by centrifugation (2,000 g, 5 min), washed twice in RPMI-1640 supplemented with 5 mM HEPES (pH 7.4) and adjusted by absorbance at 600 nm. *Candida albicans* (clinical isolate SC5314) was maintained on yeast-peptone-dextrose (YPD) agar; a single colony was grown in YPD broth for 18 h at 37 °C, sub-cultured for a further 4 h, washed twice in PBS, and quantified by absorbance at 600 nm and haemocytometer counting. Neutrophils were infected at a multiplicity of infection (MOI) of 20:1 for *S. aureus* and 5:1 for *C. albicans*.

### Induction and quantification of NET formation

Purified neutrophils were seeded at 1×10C cells per well in 96-well plates for quantification of NET formation by SYTOX Green, or in µSlide 8-well ibiTreat dishes (ibidi) for immunofluorescence and live imaging. Where indicated, cells were treated with inhibitors 30 min before stimulation. NETs were induced with phorbol 12-myristate 13-acetate (PMA, 100 nM), S. aureus (MOI 20:1) or live C. albicans (MOI 5:1). Released extracellular DNA was quantified kinetically using the cell-impermeant dye SYTOX Green (500 nM) on a [microplate reader] at 37 °C (excitation/emission [504/523 nm]) for 5 hours. For reconstitution experiments, exogenous HCOC (500 µM) or sodium hypochlorite (NaOCl, 100 µM) was added after stimulation.

### Pharmacological treatments and oxidant specificity

Inhibitor pretreatments (30 min, 37 °C) were: the MPO inhibitor 4-aminobenzoic acid hydrazide (4-ABAH, 300 µM), the NADPH-oxidase inhibitor diphenyleneiodonium (DPI, 1 µM) and the neutrophil-elastase inhibitor GW311616A (10 µM, Biomol). To probe oxidant specificity, neutrophils were treated with cumene hydroperoxide, potassium permanganate or tert-butyl hydroperoxide (10-100 µM) in parallel with HCOC (100 µM). For time-resolved inhibition, DPI was added at defined times (as indicated in the figure legend) after PMA stimulation and NET formation was quantified as above.

### Total ROS measurement

Total ROS production was measured by luminol-amplified chemiluminescence. 1×10C neutrophils were equilibrated in bicarbonate-free medium (Seahorse XF medium, Agilent) for 25 min in a COC-free incubator, pre-loaded with luminol (50 µM) with or without horseradish peroxidase (1.2 U/ml), and stimulated with PMA (50–100 nM). Luminescence was monitored for up to 3 h, and the kinetics of the oxidative burst were derived from the resulting curves.

### Detection of MPO-derived oxidants (APF) and NETotic-vacuole imaging

Intracellular hypohalous-acid production was visualized with aminophenyl fluorescein (APF, 5 µM, 5-180 min, 37 °C). Loaded neutrophils were stimulated with PMA (100 nM), *S. aureus* or live *C. albicans*; HCOC (500 µM–1 mM) served as a positive control. Kinetic fluorescence was recorded on a Fluoroskan Ascent spectrometer (excitation 488 nm, emission 550 nm), and live images were acquired on a Zeiss LSM 880 confocal microscope (excitation 488 nm; emission filter 505–550 nm). For NETotic-vacuole dynamics, time-resolved and three-dimensional (z-stack) live confocal imaging was performed at the indicated intervals over 4 h, with chromatin counterstained using Hoechst 33258 (1 µM).

### Immunofluorescence

Neutrophils were seeded on glass-bottomed dishes or coverslips, stimulated as indicated, fixed with 2% paraformaldehyde (30 min), permeabilized with 0.1–0.5% Triton X-100, and blocked (PBS with normal donkey/goat serum, cold-water-fish gelatin and BSA). Cells were stained overnight at 4 °C with primary antibodies against MPO (DAKO A0398 1:500), gp91^phox (CYBB) (sc-130543 1:200), neutrophil elastase (Cell Signalling 63610 1:1000) and p22^phox (CYBA) (sc-130551 1:200), followed by Alexa Fluor–coupled secondary antibodies (Life Technologies). DNA was counterstained with Hoechst 33258. Images were acquired on a Leica SP8 confocal microscope and colocalization was assessed in ImageJ.

### Chloride-free medium and chloride reconstitution

To assess the chloride dependence of NETosis, neutrophils were incubated in a custom chloride-free medium in which all chloride salts were replaced iso-osmotically with the corresponding gluconate salts (*37*). NET formation was quantified after PMA (100 nM) stimulation in standard RPMI, chloride-free medium, or chloride-free medium reconstituted with increasing concentrations of chloride [1-100 mM]. Osmolarity and viability controls were included. The same chloride-free and chloride-reconstituted buffers were used in the granule elastase-translocation and MPO-activity assays.

### Subcellular fractionation and isolation of granule subsets

Neutrophils (5×10C) were resuspended in disruption buffer and disrupted by nitrogen cavitation as described (*38*). Intact cells and nuclei were removed by centrifugation (300 g, 5 min). The post-nuclear supernatant, containing granules, secretory vesicles and cytosol, was loaded onto a four-layer Percoll gradient (densities 1.12, 1.09 and 1.055 g/ml, from bottom to top) and centrifuged at 37,000 g (4 °C, 20 min); twenty 2-ml fractions were collected. Fractions enriched in MPO, NGAL, MMP9 and albumin (HSA), corresponding to azurophilic, secondary (specific) and tertiary (gelatinase) granules and secretory vesicles respectively, were pooled and centrifuged at 100,000 g (4 °C, 90 min); granules and membranes were recovered as a disc above the Percoll pellet and resuspended in assay buffer. The cytosolic fraction obtained after nitrogen cavitation was retained for cytosolic neutrophil-elastase measurements.

### Immunoblotting of granule markers

Pooled granule fractions and whole-cell lysate were resolved by SDS-PAGE (equal protein per lane) and immunoblotted for compartment-specific markers: MPO (DAKO A0398 1:1000), NGAL/lipocalin-2 (secondary granules; R32169, NSJ), MMP9 (tertiary granules; clone E-11, sc-393859, Santa Cruz) and human serum albumin (secretory vesicles; MA1-19174, Invitrogen), followed by HRP-conjugated secondary antibodies; bands were developed by enhanced chemiluminescence on a Bio-Rad ChemiDoc.

### LC–MS Analysis

Samples were digested with trypsin with a filter-aided sample preparation (FASP) method as previously described (*39*). Tryptic peptides were analyzed by LC–MS using nano-flow separation on an Ultimate 3000 RSLCnano – Q-Exactive Plus (QEP) (Thermo Fisher Scientific) platform operating in data dependent acquisition mode (DDA).

### LC-MS parameters

Peptide samples were analyzed using an Ultimate 3000 RSLCnano system coupled to a QEP configured with a two-column setup. Peptides were initially concentrated on a trapping column (PepMap C18, 5 mm × 300 μm, 5 μm particle size, 100 Å pore size) for 4 min and subsequently separated on an analytical column (nano Acclaim PepMap C18, 75 μm i.d. × 250 mm, 2 μm particle size, 100 Å pore size) maintained at 40 °C. Buffer A consisted of 0.1% formic acid in water, whereas buffer B contained 80% acetonitrile with 0.1% formic acid.

FASP-digested samples were separated using an 117 min linear gradient from 3% to 55% buffer B at the same flow rate.

The QEP was dda mode with total run times of 145 min, automatically alternating between full MS and MS/MS scans. Full MS survey scans were acquired over an m/z range of 350–1650 in the Orbitrap at a resolution of 70,000 (at m/z 200). The AGC target was increased to 3 × 10C, the maximum injection time was 30 ms, and the dynamic exclusion time was 20 s. The 15 most intense multiply charged precursor ions (charge state ≥2) were selected for fragmentation by higher-energy collisional dissociation (HCD). MS/MS spectra were acquired at a resolution of 17,500 with an AGC target of 1 × 10C (FASP digestion), and maximum injection times 60 ms, respectively. A normalized collision energy (NCE) of 27% was applied for all analyses. The spray voltage was set to 2.0 kV, and the heated capillary temperature was maintained at 275 °C. Internal calibration was performed using the background ions at m/z 391.2843 and 445.1200 as lock masses.

Raw data acquired on the QEP were processed using MaxQuant (v2.0.1.0) (*40*) and searched against the reviewed human UniProt database (UP000005640, release September 2022), supplemented with common contaminant and decoy sequences. Searches were performed using trypsin/P specificity, allowing up to two missed cleavages. Carbamidomethylation of cysteine was specified as a fixed modification. For the samples, protein N-terminal acetylation and methionine oxidation were included as variable modifications.

Precursor and fragment mass tolerances were set to 10 ppm and 0.02 Da, respectively. The false discovery rate (FDR) was controlled at 1% at both the peptide-spectrum match (PSM), peptide, and protein levels.

### Scanning and transmission electron microscopy

For immunogold transmission electron microscopy (TEM), granules or cells were fixed in 2% paraformaldehyde and 0.05% glutaraldehyde, gelatin-embedded and infiltrated with 2.3 M sucrose; ultrathin cryosections were cut at −110 °C on an RMC MTX/CRX cryo-ultramicrotome, transferred to coated grids and blocked (0.3% BSA, 0.01 M glycine, 3% cold-water-fish gelatin in PBS). Sections were labelled with antibodies against MPO and neutrophil elastase, followed by donkey anti-rabbit and anti-mouse secondaries coupled to 6-nm and 12-nm gold (Jackson ImmunoResearch), contrasted with uranyl acetate/methylcellulose, and imaged on a Leo 906 TEM (Zeiss, 100 kV) with a Morada camera. For ultrastructural assessment of granule integrity, isolated granules were incubated with or without HCOC (1 or 100 µM) for 5 or 120 minutes before fixation and processing.

For scanning electron microscopy (SEM) of purified azurophilic granules, samples were fixed, mounted on stubs and coated with 3 nm platinum/carbon and imaged on LED 1550; granule diameters were measured in SmartGEM (Zeiss).

### Epoxy-ultra thin sections

For fine structural analysis, cells were fixed with 2.5% glutardialdehyde, postfixed with 0.5% osmiumtetroxide, contrasted with tannic acid and 2% uranyl acetate, dehydrated, and embedded in Polybed (Polysciences). After polymerization, specimens were cut at 60 nm and contrasted with lead citrate.

### Negative-stain Electron Microscopy

Aliquots of samples were applied to freshly glow discharged carbon-film-coated copper grids and allowed to adsorb for 10 minutes. After three washes with distilled water the grids were contrasted with 4% phospho-tungstic-acid / 1% trehalose, touched on filter paper and air-dried.

### TEM analysis

The grids were examined in a LEO 906 (Zeiss AG, Oberkochen) electron microscope operated at 100 kV and images were recorded with a Morada (SIS-Olympus, Münster) digital camera.

### Immunogold-Scanning electron microscopy

For scanning electron microscopy (SEM) of purified azurophilic granules were layered on 12 mm polylysinated coverslips, washed with PBS, blocked briefly in BSA/CWFG solution, and reacted over night with 1:1000 rb anti NE AB at 4°C. After 3 washes with PBS the samples were reacted with 18 nm-gold-conjugated goat anti rabbit IgG for 60‘, washed 4x with PBS and fixed in 2,5% glutaraldehyde. The samples were post fixed in 0,5% osmium-tetroxide, tannic acid and osmium-tetroxide again. The coverslips were then dehydrated in a graded ethanol series, dried in carbon dioxide at critical point and vacuum coated with 1.5 nm carbon-platinum. Imaging was performed using a LEO 1550 (Zeiss, Oberkochen DE) scanning-electron microscope at 20 kV acceleration voltage.

### Neutrophil elastase translocation (DQ-elastin) assay

Release of neutrophil elastase (NE) from purified azurophilic granules was measured with the membrane-impermeant fluorogenic substrate DQ-elastin (Thermofisher), whose cleavage by extragranular NE yields fluorescence. Granules were incubated in assay buffer and treated with HCOC (100 µM) in the presence or absence of 4-ABAH (300 µM); maximal release was defined by solubilization with NP-40 (0.1%) and untreated granules served as the negative control. Fluorescence was recorded kinetically (485/515 nm) for 120 minutes. Dose-response experiments used increasing HCOC concentrations. Assay specificity and pH dependence were confirmed with an alternative fluorescent streptavidin-based substrate across a range of buffer pH values as indicated in the figure legend. For genetic validation, the assay was repeated on azurophilic granules purified from MPO-deficient patients. Cytosolic NE activity was measured in the cytosolic fraction obtained by nitrogen cavitation after HCOC treatment.

### MPO enzymatic and chlorination activity

MPO chlorination activity was quantified using a taurine-chloramine assay (ab105136). Chlorination activity was assayed in parallel with the NE-translocation assay under matched chloride conditions to allow direct correlation between the two readouts.

### Chlorinating agents and lipid treatments of isolated granules

To test whether chlorinating oxidants are sufficient to mobilize NE, purified azurophilic granules were exposed to sodium hypochlorite (increasing concentrations), monochloramine (100 µM) or dichloramine (100 µM). For lipid treatments, granules or intact neutrophils were exposed to fatty acids and chlorinated fatty acids at 10 µM: palmitic acid, 2-chloropalmitic acid, stearic acid, 2-chlorostearic acid, myristic acid and arachidic acid. Lipids were prepared as stocks in ethanol or complexed with fatty-acid-free BSA; and vehicle-only controls were included throughout.

### Granule membrane-integrity (leakage) assay

Purified azurophilic granules were incubated with increasing concentrations of 2-chloropalmitic acid or with palmitic acid (control) for 120 minutes, and separated into membrane (pellet) and released (supernatant) fractions by ultracentrifugation 100xg 60 minutes). Equal-volume fractions were resolved by SDS-PAGE and stained with Coomassie; redistribution of luminal protein from the pellet to the supernatant was used as a readout of membrane permeabilization.

### Granule phospholipidomics (phosphatidylcholine and phosphatidylethanolamine)

Granule lipids were extracted in the presence of internal standards and analyzed by LC/MS as previously described (*24*, *25*, *32*). Individual choline and ethanolamine glycerophosphospholipids were detected using parallel reaction monitoring with an Q-Exactive mass spectrometer equipped with a Vanquish UHPLC System (Thermo Scientific) with isotopomer corrections for each target molecular species compared to the respective internal standard. Lipids were separated by reversed phase chromatography with an AccucoreTM C30 column 2.1 mm × 150 mm (Thermo Scientific) stationary phase and a gradient of mobile phases comprised of either 60% acetonitrile, 40% water, 10 mM ammonium formate, and 0.1% formic acid (mobile phase A) or 90% isopropanol, 10% acetonitrile with 2 mM ammonium formate, and 0.02% formic acid (mobile phase B).

### Quantification of chlorinated lipids (2-chlorofatty aldehydes and 2-chlorofatty acids)

Granule 2-chlorofatty aldehydes or 2-chlorofatty acids were extracted in the presence of internal standards and analyzed by GC/MS or LC/MS, respectively, as previously described (*25*, *30*, *31*). Prior to GC/MS 2-chlorofatty aldehydes were converted to their pentafluorobenzyl oxime derivatives and were detected by negative ion chemical ionization detection. Internal standards were 2-chloro-[7,7,8,8-d4]-hexadecanal and 2-chloro-[7,7,8,8-d4]-hexadecanoic acid for quantification of 2-chlorofatty aldehydes and 2-chlorofatty acids, respectively.

### Statistical analysis

Statistical analyses were performed in GraphPad Prism v10 and the R statistical environment. Data are presented as [mean ± s.d. or s.e.m.], with individual data points shown where applicable; n denotes the number of independent biological replicates (donors or granule preparations) and is stated in each figure legend. Specific tests are indicated per figure (e.g. paired two-tailed Student’s t-test for before/after granule measurements; one-way or two-way ANOVA with Dunnett’s or Tukey’s multiple-comparison correction for multi-group analyses). Exact P values are reported in the figures, and differences were considered significant at P < 0.05.

## Data availability

The datasets generated and/or analyzed during the current study, including raw and processed data, are available from the corresponding authors upon reasonable request. Any additional materials, protocols, or code used in the study are also available upon request to ensure reproducibility of the results.

## Supplementary figures

**Figure S1.**
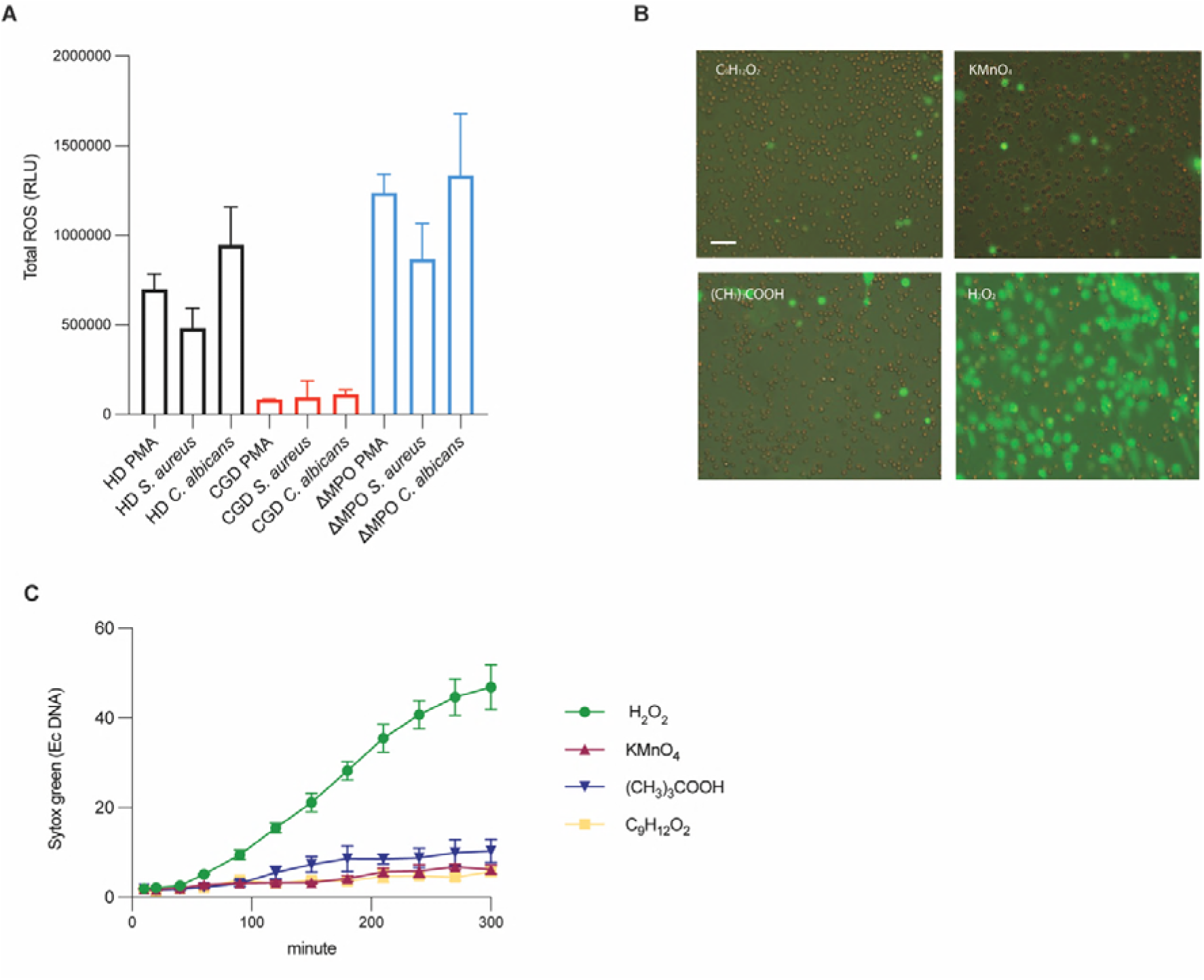
NET formation depends on MPO-derived oxidants. (A) Total reactive oxygen species (ROS) production in neutrophils from healthy donors, CGD patients, and MPO-deficient patients stimulated with PMA or C. albicans for 120 min (N as indicated). (B) Representative live-cell epifluorescence microscopy images of neutrophils from healthy donors stimulated for 5 h with cumene hydroperoxide (100 μM), potassium permanganate (100 μM), tert-butyl hydroperoxide (100 μM), or hydrogen peroxide (100 μM) (N = 3). Draq5 stains intracellular DNA, and Sytox Green labels extracellular DNA. Scale bar, 50 μm. (C) Kinetics of NET formation in neutrophils from healthy donors stimulated with cumene hydroperoxide (100 μM), potassium permanganate (100 μM), tert-butyl hydroperoxide (100 μM), or hydrogen peroxide (100 μM), quantified over time based on Sytox Green fluorescence (N = 3).

**Figure S2.**
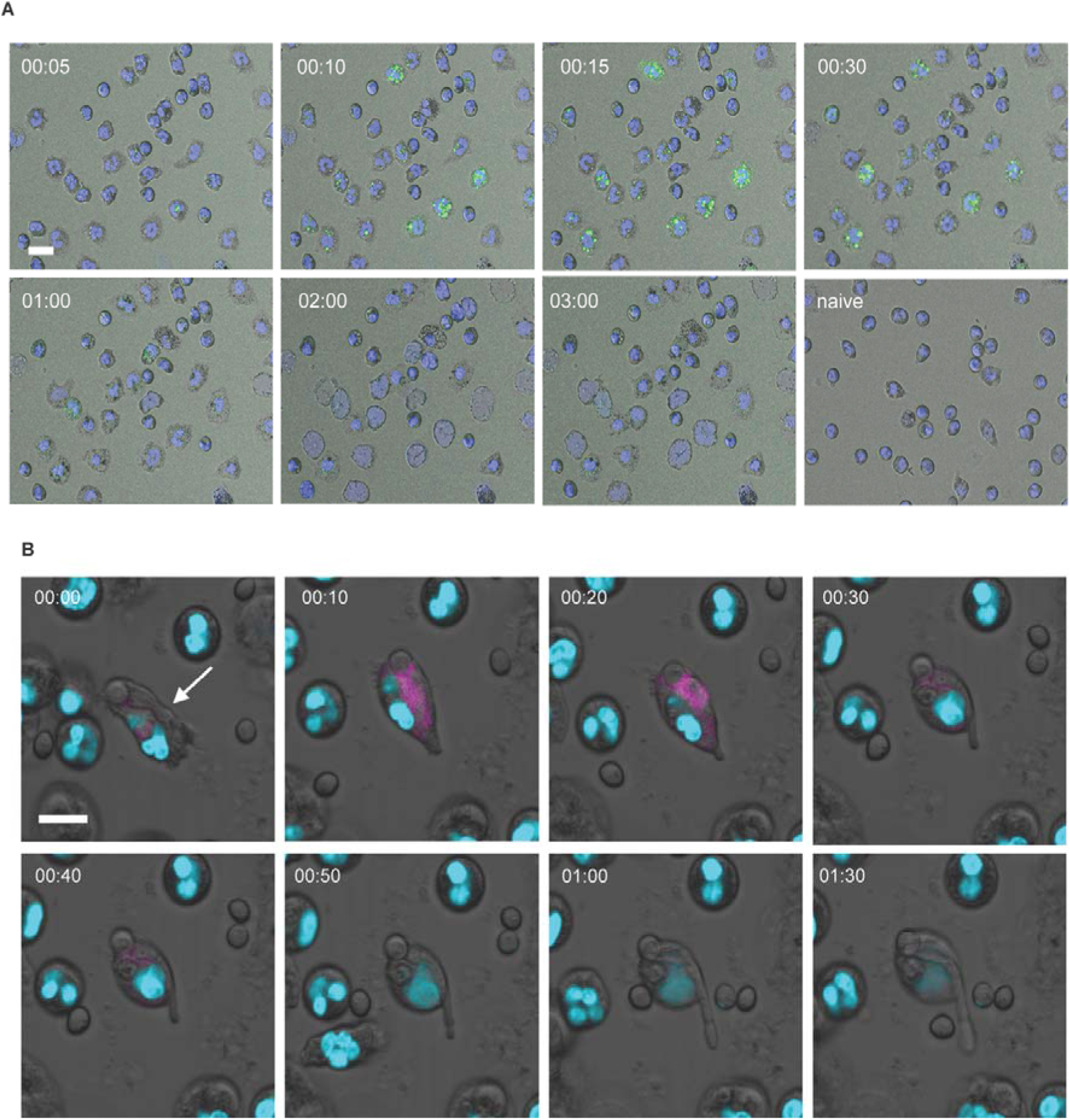
Intracellular dynamics of MPO-derived oxidants during NETosis. (A) Time-course live confocal microscopy images of neutrophils from a healthy donor stimulated with PMA (100 nM) and imaged at 5, 10, 15, 30, 60, 120, and 180 min after stimulation, or left unstimulated (N = 5). Blue fluorescence indicates intracellular DNA, and green fluorescence indicates intracellular hypohalous acid formation. Scale bar, 10 μm. (B) Time-course three-dimensional live confocal microscopy images of neutrophils from healthy donors stimulated with live *C. albicans* (MOI 5:1) and imaged at 0, 10, 20, 30, 40, 50, 60, and 90 min after stimulation (N = 3). Light blue fluorescence indicates intracellular DNA, and magenta fluorescence indicates intracellular hypohalous acid formation. Scale bar, 10 μm.

**Figure S3.**
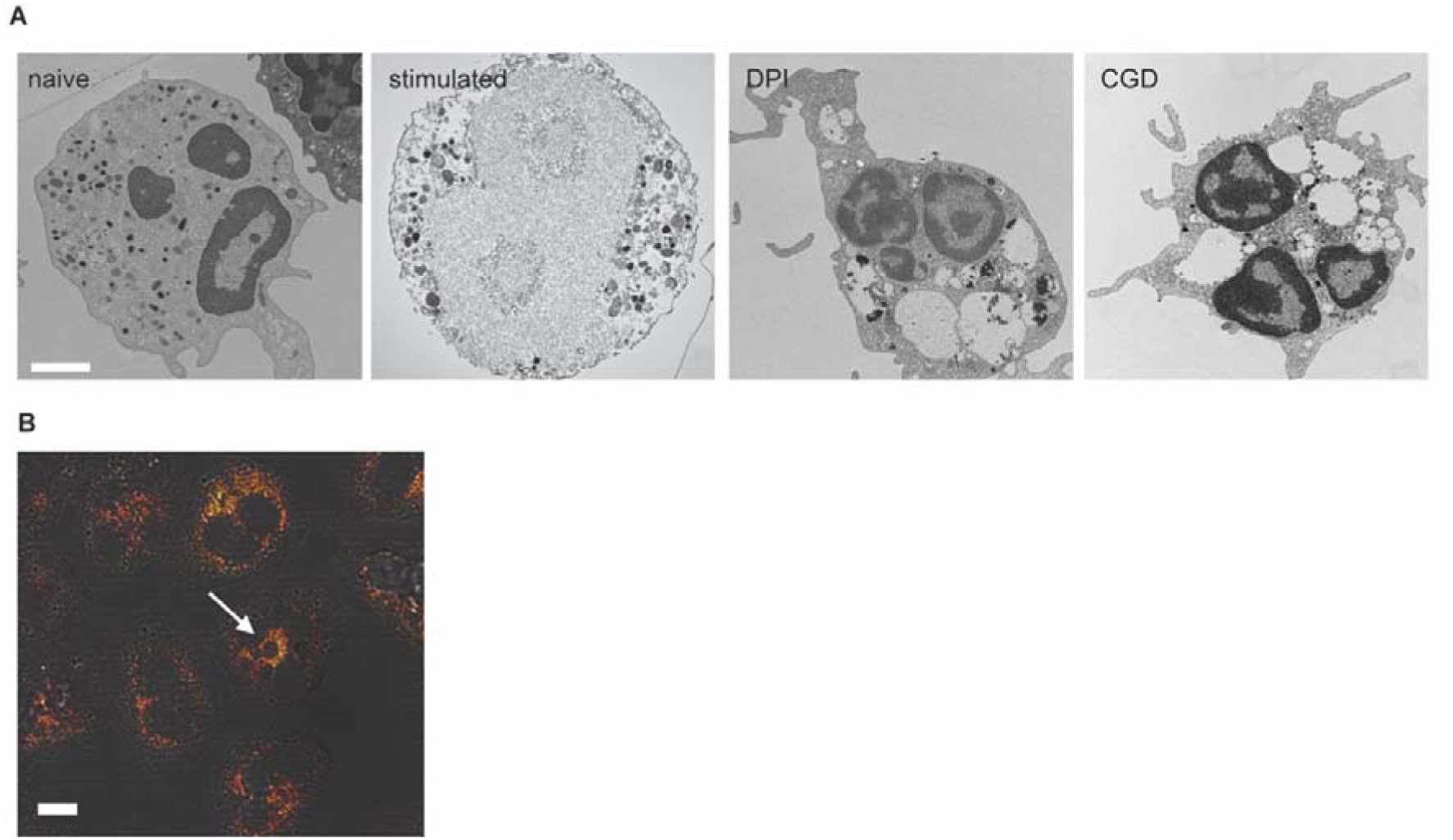
ROS are required for NETotic vacuoles ultrastructural changes and granule protein association with the plasma membrane. (A) Transmission electron microscopy images of healthy donor neutrophils left untreated (naïve), stimulated with PMA for 3 h, pretreated with DPI before PMA stimulation, or CGD neutrophils stimulated with PMA (N = 3). Scale bar, 0.5 μm. (B) Confocal immunofluorescence images of healthy donor neutrophils stimulated with PMA for 30 min and stained for neutrophil elastase (NE) and p22^phox^ (N = 3). Scale bar, 2.5 μm.

**Figure S4.**
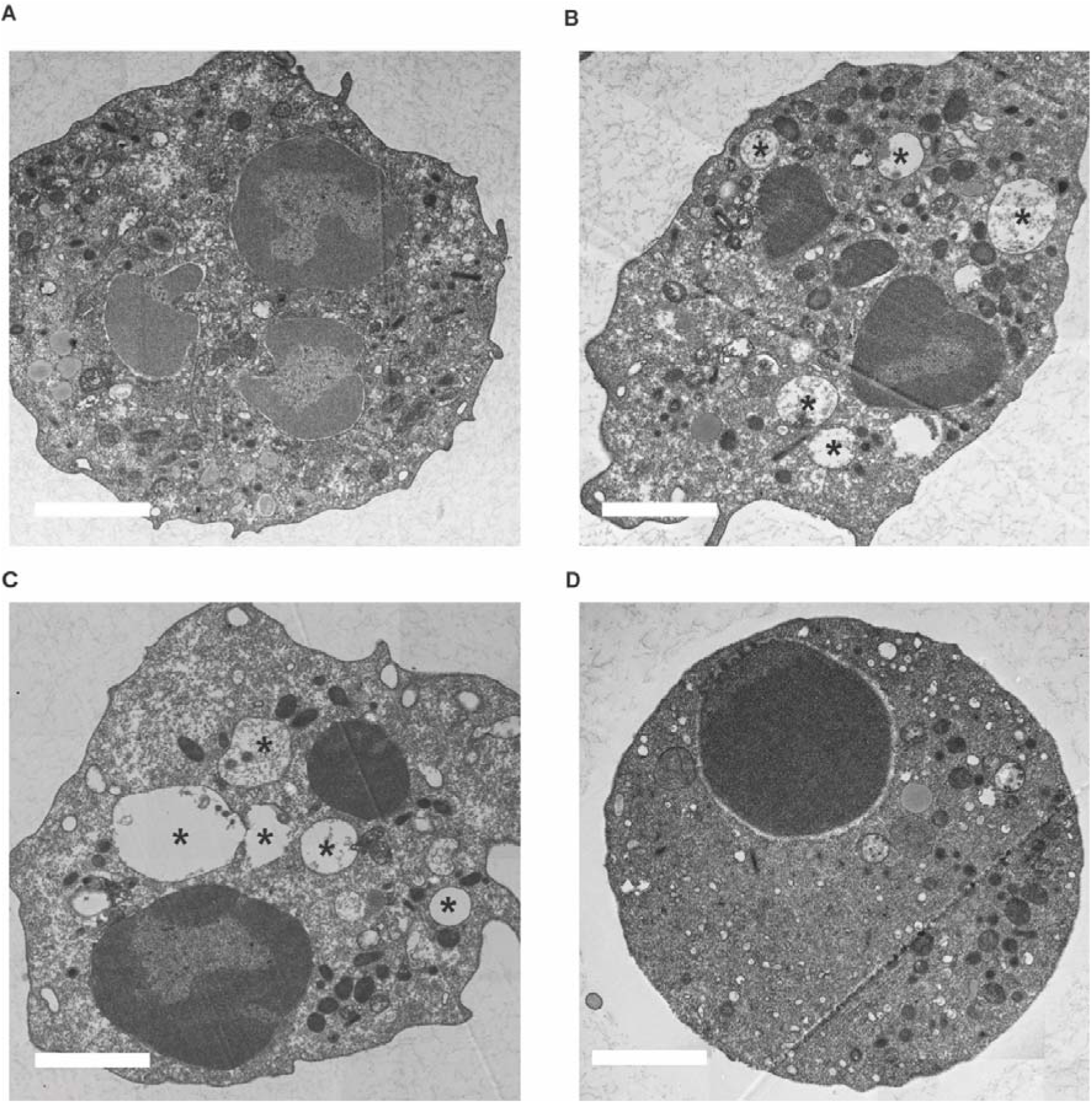
Ultrastructural time course of NETotic vacuole formation during PMA-induced NETosis. Transmission electron microscopy images of a healthy donor neutrophil at baseline and following stimulation with PMA (100 nM) for the indicated times. (A) 0 min (naïve/unstimulated), (B) 4 min, (C) 10 min, and (D) 60 min after PMA stimulation. Asterisks (*) indicate NETotic vacuoles. Scale bar, 0.5 μm.

**Figure S5.**
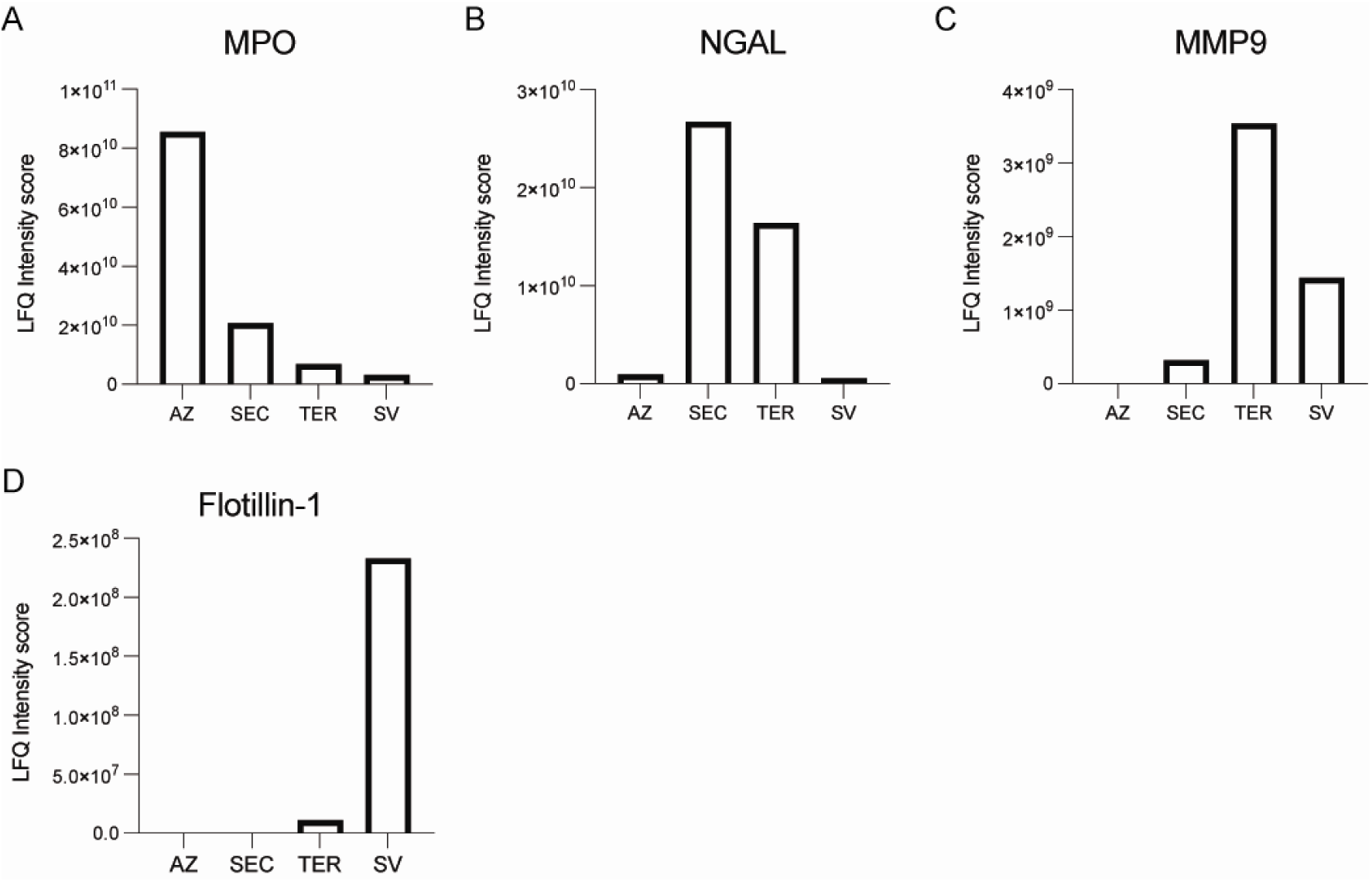
Proteomic validation of neutrophil granule subset purity. Label-free quantification (LFQ) intensity of established granule-marker proteins across four purified granule fractions — azurophilic (AZ), secondary/specific (SEC), tertiary/gelatinase (TER) granules and secretory vesicles (SV) — separated by Percoll density-gradient fractionation and analysed by liquid chromatography–tandem mass spectrometry. (A) Myeloperoxidase (MPO), a marker of azurophilic granules, is most abundant in the AZ fraction. Bars represent the mean of three technical replicates from one donor, representative of three independent donors. (B) Neutrophil gelatinase-associated lipocalin (NGAL/lipocalin-2), a marker of secondary granules, is enriched in the SEC fraction. Bars represent the mean of three technical replicates from one donor, representative of three independent donors. (C) Matrix metalloproteinase-9 (MMP9), a marker of tertiary granules, is enriched in the TER fraction. Bars represent the mean of three technical replicates from one donor, representative of three independent donors. (D) Flotillin-1, a marker of secretory vesicles, is enriched in the SV fraction. Bars represent the mean of three technical replicates from one donor, representative of three independent donors.

**Figure S6.**
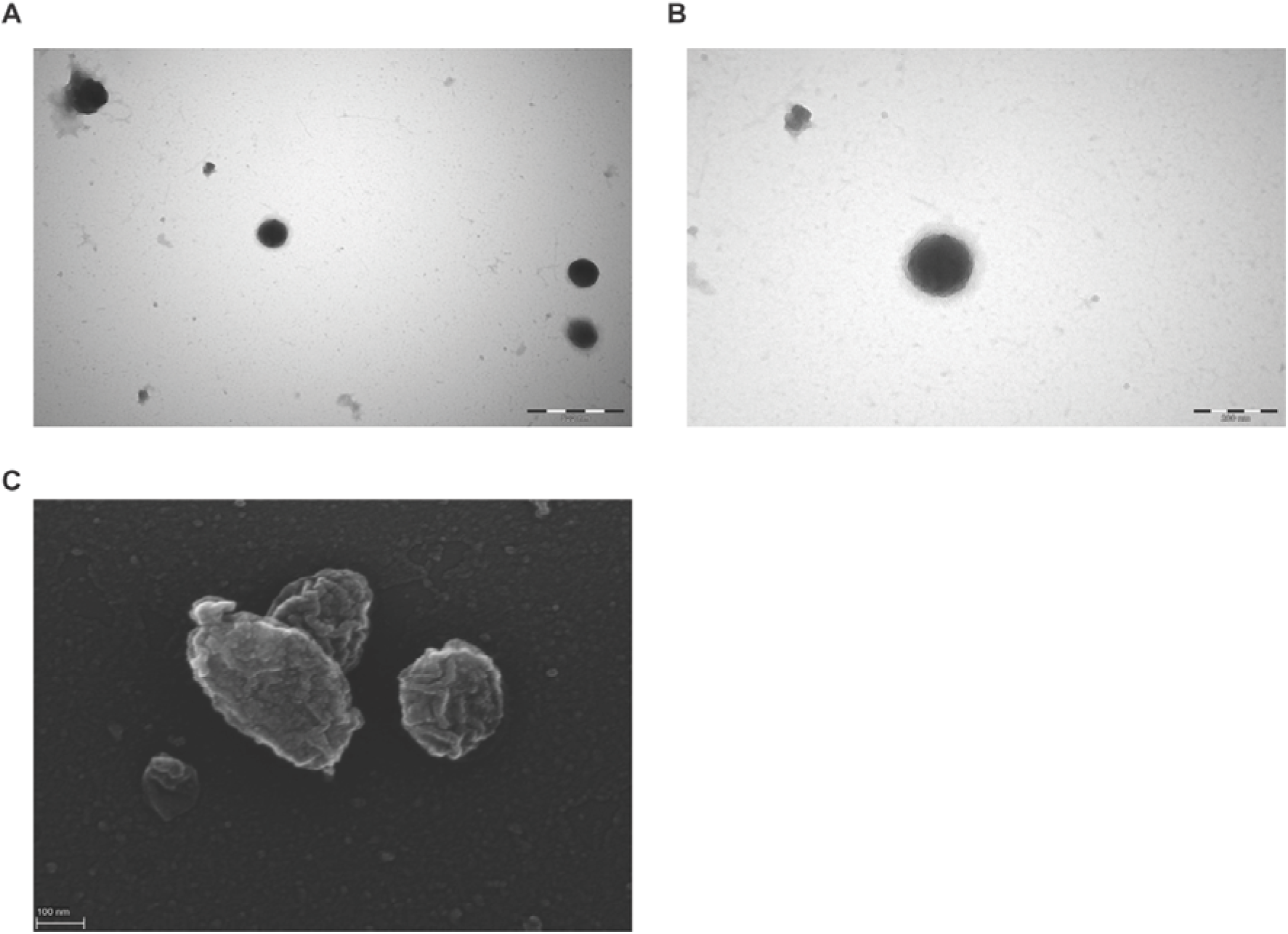
Ultrastructure of azurophilic granules. (A–B) Negative-stain electron microscopy images of azurophilic granules purified from healthy donor neutrophils (N = 4). Scale bar, 200 nm. (C) Scanning electron microscopy images of purified azurophilic granules either unstained (N = 4).

**Figure S7.**
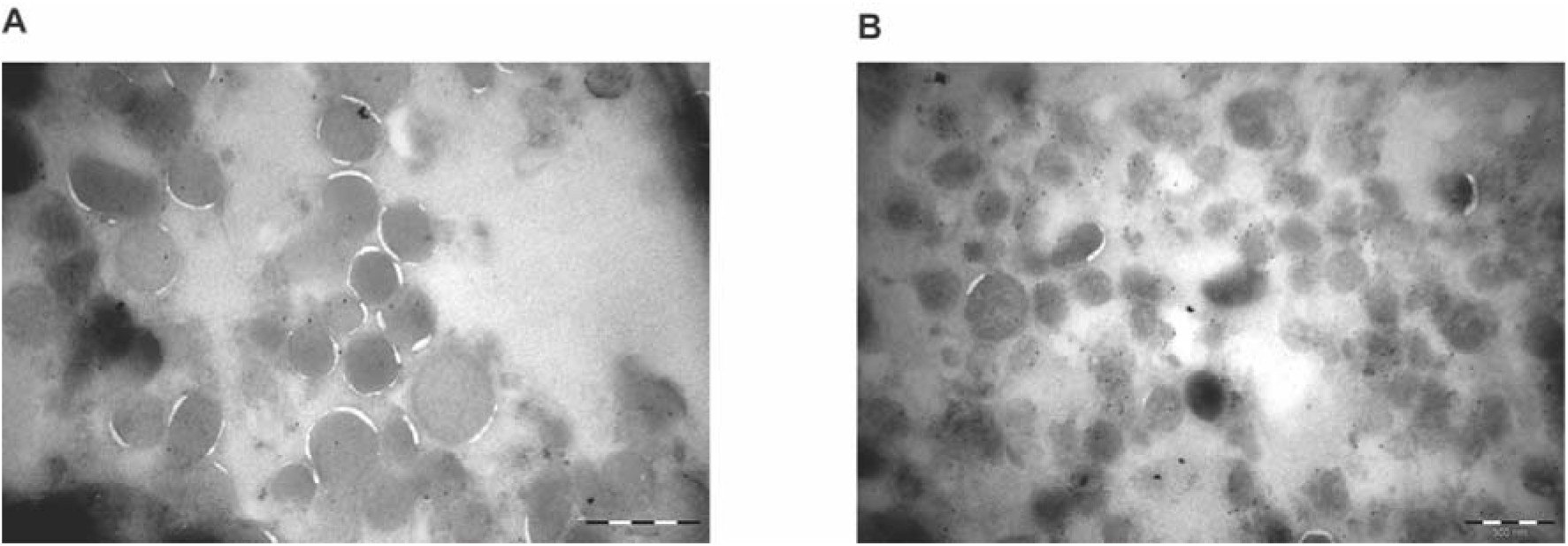
Azurophilic granules stimulated with exogenous hydrogen peroxide. (E–F) Transmission electron microscopy images of azurophilic granules either treated with H□O□ 1 μM for 5 minutes (A) or 100 μM for 2 h (N = 3).

**Figure S8.**
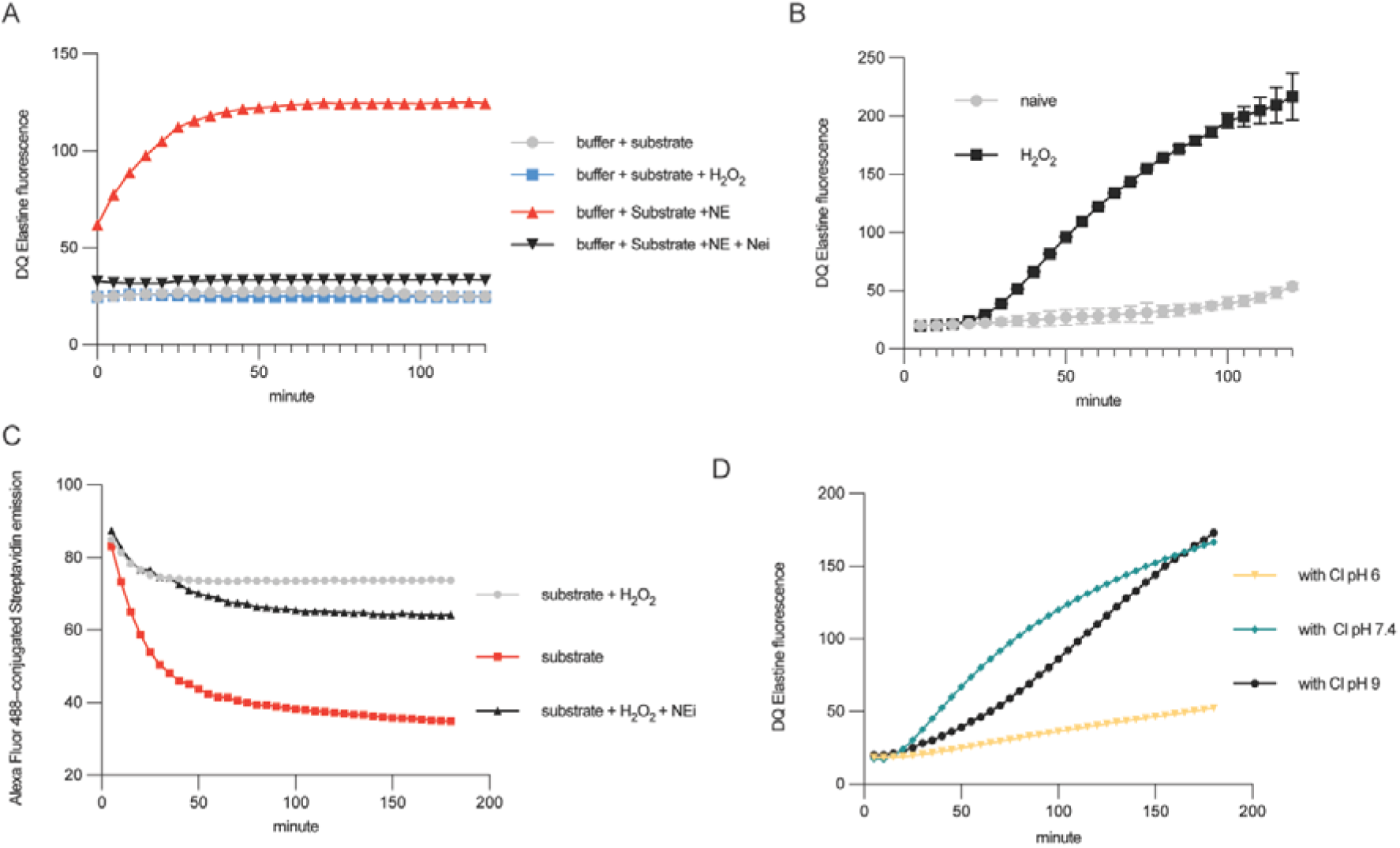
Dynamic mobilization of neutrophil elastase from azurophilic granules. (A) Elastin degradation assay measuring neutrophil elastase (NE) activity in purified azurophilic granules using a fluorescent elastin substrate. Fluorescence increase over time reflects NE-mediated elastin cleavage (N = 3). (B) NE translocation assay in cytosolic fractions of neutrophils isolated by nitrogen cavitation. Cytosolic NE activity was measured after stimulation with H□O□, indicating release of NE from azurophilic granules into the cytosol (N = 3). (C) NE translocation assay using a fluorescent streptavidin-based substrate to detect NE activity in cytosolic fractions following H□O□ stimulation (N = 3). (D) NE translocation assay performed at different pH conditions after H□O□ treatment to evaluate the effect of pH on NE mobilization and activity (N = 3).

**Figure S9.**
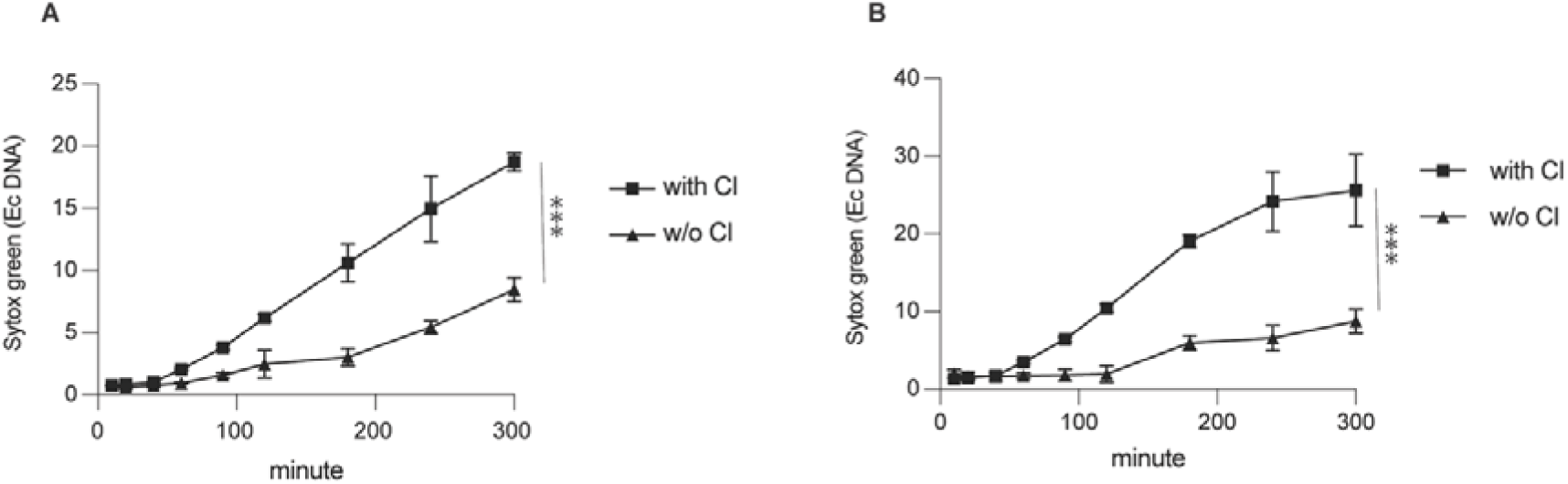
Chloride dependence of pathogen-induced NET formation. Time-course quantification of NET formation, measured by Sytox Green fluorescence (extracellular DNA), in neutrophils from healthy donors stimulated with (A) *Staphylococcus aureus* or (B) *Candida albicans* in the presence (“with Cl□”) or absence (“w/o Cl□”) of chloride. Data are mean ± SEM. ***P < 0.001.

**Figure S10.**
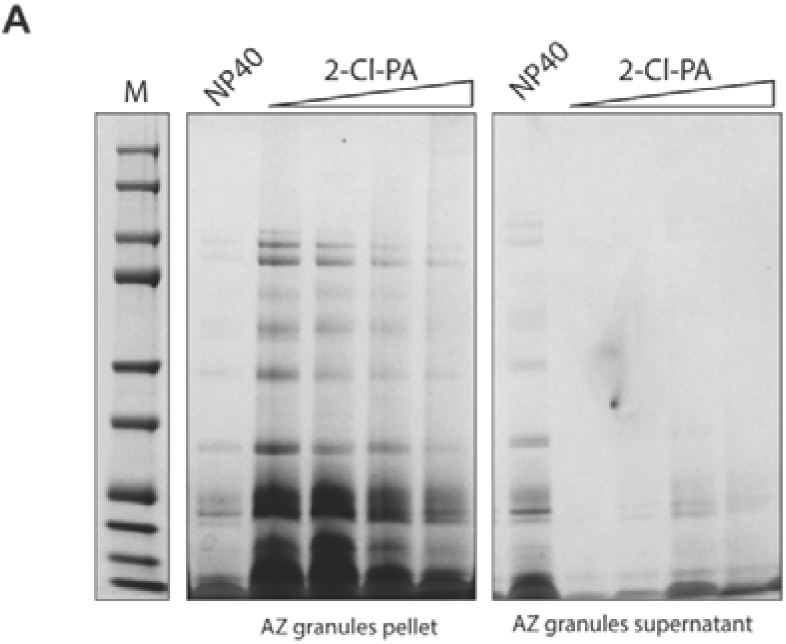
2-chloropalmitic acid induces protein release from azurophilic granules. (A) SDS-PAGE analysis of azurophilic (AZ) granules treated with NP-40 (positive control, total lysis) or increasing concentrations of 2-chloropalmitic acid (2-Cl-PA), followed by ultracentrifugation. Total protein content is shown in the granule pellet (left) and supernatant (right) fractions. M, molecular weight marker.

